# Contractile function maintains cardiomyocyte differentiation and inhibits cell cycle activity

**DOI:** 10.1101/2025.08.19.671044

**Authors:** Lynn A.C. Devilée, Janice D. Reid, James R. Krycer, Harley R. Robinson, Sean J. Humphrey, Chris Siu Yeung Chow, Mary Lor, Rebecca L. Fitzsimmons, Qing-Dong Wang, Michael Doran, Enzo R. Porrello, Nathan J. Palpant, Simon R. Foster, Richard J. Mills, James E. Hudson

**Affiliations:** Cardiac Bioengineering Laboratory, QIMR Berghofer, Brisbane, QLD, Australia; Cardiac Drug Discovery Laboratory, QIMR Berghofer, Brisbane, QLD, Australia; Mucosal Immunology Laboratory, QIMR Berghofer, Brisbane, QLD, Australia; Faculty of Health, School of Biomedical Sciences, Queensland University of Technology, Brisbane, QLD, Australia; Functional Phosphoproteomics, Murdoch Children’s Research Institute, Royal Children’s Hospital, Melbourne, VIC, Australia; Institute for Molecular Bioscience, University of Queensland, QLD, Australia; Department of Pharmacology, Monash Biomedicine Discovery Institute, Monash University, Melbourne, VIC, Australia; Synthetic Cell Therapeutics, LLC, Gaithersburg, Maryland, USA; Murdoch Children’s Research Institute, Royal Children’s Hospital, Melbourne, Victoria, Australia; Novo Nordisk Foundation Centre for Stem Cell Medicine (reNEW), Murdoch Children’s Research Institute, Melbourne, Victoria, Australia; Department of Paediatrics, School of Biomedical Sciences, University of Melbourne, Melbourne, Victoria, Australia; Department of Anatomy & Physiology, School of Biomedical Sciences, The University of Melbourne, Melbourne, Victoria, Australia; Melbourne Centre for Cardiovascular Genomics and Regenerative Medicine, Melbourne, Victoria, Australia; School of Biomedical Sciences, University of Queensland, QLD, Australia

## Abstract

Numerous endotherm species lose cardiac regenerative capacity shortly after birth, which is in contrast to many ectotherm species who regenerate throughout life. Whether the enhanced contractile function required for endothermy contributes to the cell-cycle exit remains to be explored. Herein, we use human cardiac organoids with advanced maturation combined with direct targeting of contraction using mavacamten and aficamten to enable exquisite control of active contraction over brief time windows. We show that transient inhibition of contraction re-activates the cell cycle. Multi-omics analyses demonstrated the cell cycle response to be mediated through a dedifferentiation-like process, which was swiftly reversed upon removal of the myosin inhibitors. Together these findings reveal that active contraction maintains differentiation including cell cycle arrest in cardiomyocytes.

## Introduction

The turnover rate of adult human cardiomyocytes has been estimated to be <1% per year, a rate which precludes the repair of damage due to a myocardial infarction or chronic insults (1, 2). However, during a brief neonatal window, the human heart is able to restore functionality following damage as discussed in a handful of case reports (3–7). While cardiomyocyte regeneration is only inferred in these cases, based on studies in neonatal mice which have regenerative capacity during the first week after birth, it is likely that cardiomyocyte regeneration via proliferation contributed to recovery (8). The brief neonatal regenerative time-window is present in different endotherm (warm-blooded) species such as pigs, rabbits, rats and sheep (9, 10), while many ectotherms regenerate throughout life (9, 11–13). Hence, reactivating the cell cycle of mature cardiomyocytes offers a potential avenue to transform heart failure treatment.

The decline in cardiomyocyte proliferative capacity coincides with an extensive maturation process of cardiomyocytes, which shifts their metabolic profile towards oxidative metabolism, improves their electrophysiological properties and enhances their contractile power and efficiency (14). Adult cardiomyocytes differ in many aspects from their immature counterparts, with over 6000 differentially expressed genes between the two states (15). To regenerate, zebrafish and neonatal mouse cardiomyocytes undergo (transcriptional) dedifferentiation, yet in response to damage, adult mouse cardiomyocytes fail to revert to such immature states (11, 15, 16). Attempts to reverse the mature phenotype through forced dedifferentiation induced by overexpression of the Yamanaka factors Oct4, Sox2, Klf4, and C-Myc (OSKM) (17–20), constitutively active Yap (21, 22) or Erbb2 (23, 24) effectively induced cardiomyocyte proliferation *in vitro* and *in vivo*. Thus, the advanced level of maturity of adult cardiomyocytes may pose a barrier to regeneration.

A collection of literature has provided evidence demonstrating that forcing mature cardiomyocytes to proliferate decreases their active force production. Ejection fraction or contraction force was reduced in a long-term OSKM overexpression model (19), as a consequence of constitutively active Erbb2 in rat and human cardiomyocytes (24), in mature human cardiac organoids (hCOs) treated with the GSK-3α/β inhibitor CHIR99021 (25) and in a pro-proliferative compound screen in hCOs (26). Likewise, induction of cytokinesis through overexpression of Plk1 and Ect2 came at the expense of contraction in adult mice (27) while knockdown of *Uqcrfs1*, a component of complex III of the electron transport chain, activated cardiomyocyte proliferation in mice by day 60, which coincided with a reduction in ejection fraction (28). Compound heterozygous *ALMS1* mutations encoding Alström syndrome protein 1 in humans and mice also led to reduced contractile function and prolonged cell cycle activity after birth, eventually resulting in cardiomyopathy (29, 30). The potential involvement of cardiac contraction in regulating cardiac proliferation is moreover depicted by the segregation of regenerative capacity between regenerative ectotherms and non-regenerative endotherms (9, 10). Endotherm species, including Tuna fish which display regional endothermy of the brain and eyes (31), and the only fully endotherm fish known to date, the small eye opah (Lampris spp) (32, 33), have a stronger cardiovascular function compared to ectotherms which is in part mediated by the neonatal maturation phase (14, 34). Impaired cardiovascular responsiveness during cold exposure elevates the risk of hypothermia, underscoring the importance of robust cardiac function for maintaining endothermy (35, 36). These different studies may converge on a mechanism where the need for strong cardiac function blocks cell cycle re-entry of mature endotherm cardiomyocytes, captured in the statement of Rumyantsev in 1984 (37), “cardiomyocytes, being *sui generis* the slaves of their uninterrupted function, have received no less heavy fetters in adapting themselves to reproduction in the differentiated state”.

Recently, we and others have explored more direct and precise interrogation of the role of cardiac function in blocking cardiomyocyte proliferation. Inhibition of the L-type calcium channels (LTCC), which consequently disrupts excitation-contraction coupling, was demonstrated to be pro-proliferative in 2D cardiomyocytes, hCOs and mouse models (38, 39). Slowing down the heart rate in adult mice also successfully led to proliferation, and there were signs of cardiac proliferation in human hearts following mechanical unloading after left ventricular assist device implantation (40–42). However, interference through perturbed calcium cycling or mechanical unloading impacts many critical cardiomyocyte functions and it is challenging to dissect which processes are critical for the proliferative response.

Herein, we assess the direct impact of cardiac contractility on cell cycle activity. Using direct inhibition of active force production with cardiac myosin modulators, we demonstrate that this re-activates cardiomyocyte cell cycle in hCOs. A multi-omics approach revealed that cardiac contractility directly imposes a mature cardiomyocyte state. When contraction is temporarily interrupted, a dedifferentiation-like response allows for endogenous cell cycle reactivation.

## Results

### Myosin super relaxed state stabilization reduces active force and induces cell cycle activity in hCOs

Mavacamten was recently approved for the treatment of hypertrophic cardiomyopathy, where it is used to reduce contractility and energy demand by stabilizing the myosin heavy chain in an energy-conserving ‘super relaxed state’ (43, 44). Here we used mavacamten in healthy hCOs to assess how contractility impacts cardiac cell cycle activity (Figure 1a-b). Mavacamten significantly reduced active force of contraction within 1 h of treatment, which was maintained throughout 48 h (Figure 1c-d). At concentrations that did not completely abolish contractile function (0.1 and 0.3 µM), contraction rate as well as activation and relaxation kinetics were not impacted (Figure 1c). Additionally, mavacamten did not suppress calcium transients (Figure 1e), highlighting the specificity of mavacamten for altering myosin-controlled active force.

**Figure 1.**
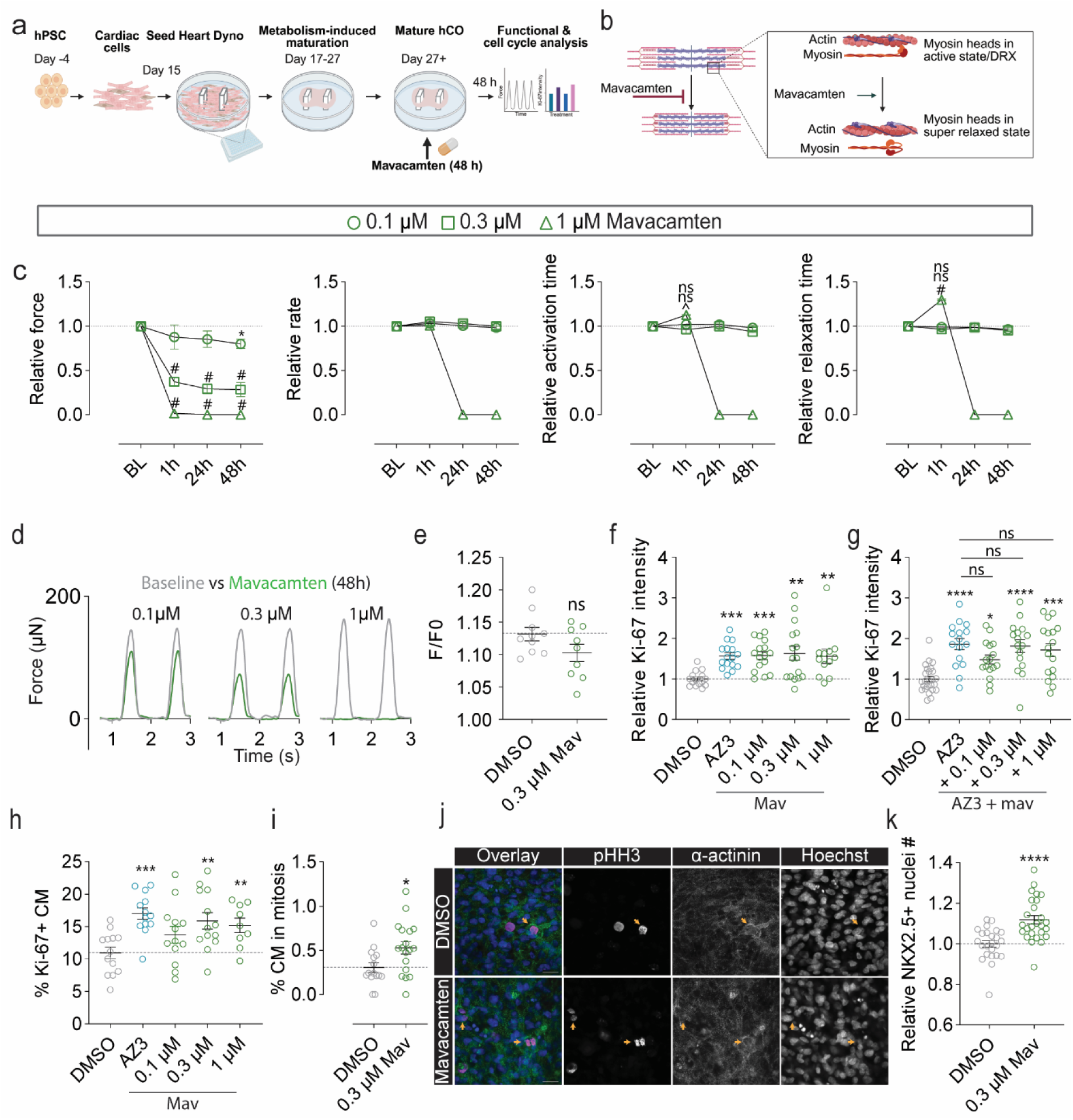
Mavacamten initiates cell cycle activity in mature hCOs. (a) Experimental design. hCOs were treated for 48 h with AZ3 (3 µM) or 0.1, 0.3 or 1 µM mavacamten. (b) Schematic of the mechanism of action of mavacamten, displaying how mavacamten stabilizes the myosin heads in the super relaxed state. (c) Assessment of functional hCO parameters over time relative to baseline normalized to control. Mavacamten concentration-dependently reduces force of contraction without affecting rate, 50% activation time or 50% relaxation time at 0.1 or 0.3 µM. N = 19-32 hCOs from 2-3 experiments. (d) Representative force curves for an individual hCO before (baseline) and after 48 h of mavacamten treatment. (e) Mavacamten did not significantly impact Ca^2+^ release as determined by Fluo-4 immunofluorescent intensity. N = 9-10 hCOs from 2 experiments. (f) Quantification of whole hCO Ki-67 intensity suggests cell cycle activation induced by mavacamten. N = 12-16 hCOs from 2-3 experiments. (g) Combination of AZ3 with mavacamten did not boost overall levels of Ki-67 whole tissue intensity. AZ3 was used at 3 µM. N = 16-25 hCOs from 2 experiments. (h) Manual quantification of the percentage of cardiomyocytes positive for Ki-67 demonstrated 0.3 µM mavacamten was most effective in initiating cell cycle activity. N = 9-13 hCOs from 2-3 experiments. (i) Quantification of the percentage of cardiomyocytes in mitosis based on pHH3 expression and sarcomere remodeling. 0.3 µM effectively increased cardiac mitosis. N = 15-18 hCOs from 4 experiments. (j) Representative images of cardiomyocytes in late mitosis stages as marked by pHH3 expression and actin remodeling (arrowheads). pHH3 in magenta, Hoechst in blue, α-actinin in green. Scale bar = 20 µm. (k) Automated analysis of the number of cardiomyocyte nuclei based on NKX2-5 expression highlights mavacamten supported karyokinesis completion after 48 h. N = 24-26 hCOs from 3 experiments. Due to the nature of the original screen with additional compounds, some control data points overlap with data published in Devilée et al. (Figure 1b, d, e and figure S1) (39). *p < 0.05, ^/**p < 0.01, ***p < 0.001, #/****p < 0.0001 using a Two-way ANOVA with Dunnett’s test for multiple comparisons (c), a Kruskal-Wallis test with Dunn’s test for multiple comparisons (f), one-way ANOVA with Dunnett’s test for multiple comparisons (g,h), or an unpaired t-test (e, i, k). Significance is depicted top to bottom for 0.1, 0.3 and 1 µM mavacamten, respectively (c). Data are presented as mean ± SEM with individual data points representing average of all experiments (c) or individual hCOs (e-k). Experiments were performed using the HES3 cell line cultured using the FBS media protocol (25, 68). *BL baseline, Mav mavacamten*.

We next assessed whether contraction controls cell cycle activity and used AZ3, a pro-proliferative compound, as a positive control (26). Analysis of cell cycle activation demonstrated an increase in whole hCO Ki-67 intensity across all three mavacamten concentrations (Figure 1f). An additive effect of AZ3 and mavacamten on cell cycle activation was not observed (Figure 1g), indicating that these treatments may share pathways for activation. The mavacamten proliferation response plateaued at 0.3 µM for the percentage of cardiomyocyte nuclei positive for Ki-67 (Figure 1h) and at this concentration, cardiomyocytes progressed to mitosis (Figure 1i-j) and karyokinesis with an 11% increase in cardiomyocyte nuclei number (Figure 1k) assessed by automated quantification of whole hCO (Figure S1). The robustness of the response was confirmed using aficamten, a myosin inhibitor that binds to a distinct site on the myosin heavy chain, yet with similar functional consequences (45). Aficamten treatment for 48 h reduced active force production while also stimulating cardiac cell cycle activity across a range of concentrations (Figure S2). These findings corroborate with the level of cell cycle activity when the LTCC is inhibited in hCOs (39), indicating that this may in-part rely on reducing active force.

We next assessed whether mavacamten-induced cell cycle activity is enhanced by co-treatment with FGF2, a growth factor known to enhance cardiomyocyte proliferation during development and an important protective mediator of the injury response (46, 47). FGF2 alone induced an increasing trend in Ki-67 positive cardiomyocyte nuclei, which may be due to activation of the initial cell cycle stages (hypertrophy) (48). Combination treatment robustly increased overall cell cycle activity yet failed to boost cardiomyocyte nuclei number (Figure S3a-d), despite more cardiomyocytes in mitosis compared to hCOs treated with mavacamten alone (Figure S3e). Hence, cardiomyocytes may exit the cell cycle prior to karyokinesis or die shortly after, potentially related to the rescue of cardiac contractility by FGF2.

### Cardiomyocyte cell cycle induction is transient

To determine whether the cell cycle is activated by an abrupt change or sustained over a longer period, hCOs were treated for 168 h (Continuous, Con), for 48 h and then recovered for 120 h (Recover, Rec) or for 48 h, recovered for 72 h and then treated again for 48 h (Repeat, Rep) (Figure 2a). Here, AZ3 and nifedipine were included as control groups. The functional parameters followed the expected patterns for each treatment regimen (Figure 2a-d, Figure S4). In hCOs treated continuously with AZ3, nifedipine or mavacamten, levels of Ki-67 returned to baseline and there was no further boost in cardiomyocyte nuclei number, indicating a transient induction of the cell cycle (Figure 2e-f). Cell cycle activity returned to baseline in the recovery groups while it was re-induced in the repeat groups (Figure 2e). There was a mild drop in cardiomyocyte nuclei number in the control hCOs at experimental end point, which is likely the direct result of long-term culture. A similar decrease was observed in mavacamten-treated hCOs. Continuous treatment with nifedipine however resulted in a further reduction in cardiomyocyte nuclei number by 168 h, indicating that long-term inhibition of calcium cycling is detrimental whereas transient treatment in the recovery and repeat groups maintained the increased nuclei number (Figure 2f). The increase in cardiomyocyte nuclei was lost in the recovery group with AZ3 treatment, which may indicate that the cardiomyocytes become addicted to AZ3 and removal causes cell stress and death (Figure 2c, f). Importantly, the number of cardiomyocyte nuclei in the mavacamten-recovery group was maintained and function fully recovered, even exceeding baseline force by 10%, which indicates the 11% additional cardiomyocytes contribute to increased contractile force (Figure 2c, f). Collectively, inhibition of contractility results in a transient induction of cardiomyocyte cell cycle activity, the results of which are still present after removal of mavacamten and full force recovery.

**Figure 2.**
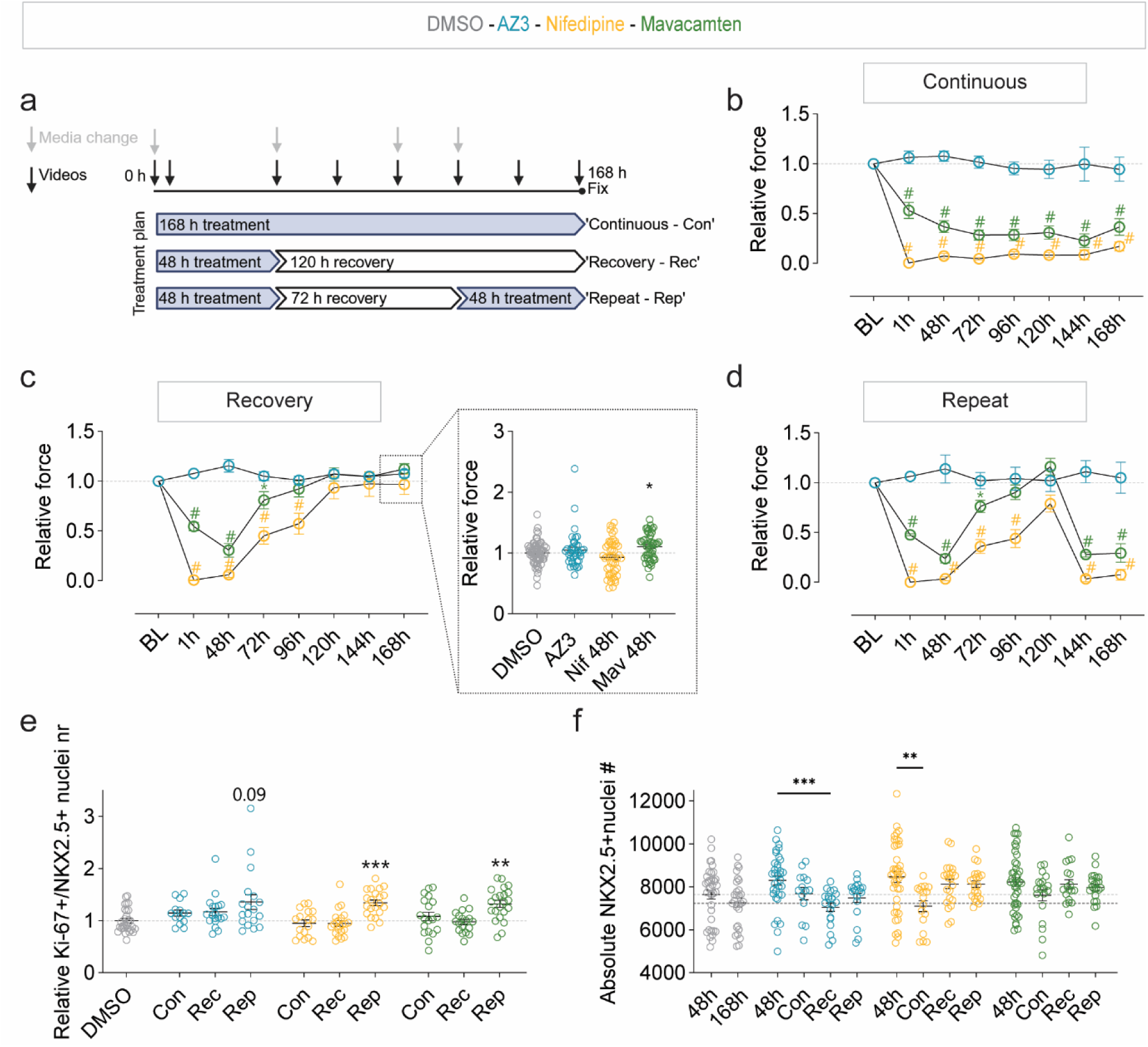
Cardiomyocytes transiently re-enter the cell cycle. (a) Schematic of experiment where hCOs were treated for 168 h (continuous), for 48 h with 120 h recovery (recovery) or repeatedly for 48 h with 72 h no treatment in between treatment periods (repeat). AZ3 was used at 3 µM, nifedipine at 3 µM and mavacamten at 0.3 µM. (b) Force of contraction during 168 h of continuous treatment with AZ3, nifedipine or mavacamten. N = 4-5 experiments. (c) Assessment of hCO functional recovery after ceasing AZ3, nifedipine or mavacamten treatment. Force returns to baseline levels. Inset displays hCO force at endpoint (168 h) highlighting hCOs treated with mavacamten displayed a slight increase in contractile force. N = 5-6 experiments. (d) Force of contraction after repeated treatments shows a consistent functional response to nifedipine and mavacamten with sufficient time in between treatment periods to fully recover function. N=4 experiments. (e) Overall cell cycle activity across the different treatment time frames. N = 15-33 hCOs from 4-6 experiments. (f) Absolute cardiomyocyte nuclei number in one z-plane at experimental endpoints with 48 h being 48 h experiments, while all other conditions relate to the long-term experiments as described in (A). The grey dashed line indicates the average nuclei number after 48 h experiments, the black dashed line indicates the average nuclei number after 168 h experiments. N = 14-29 hCOs from 4 experiments (7-day experiments) and N = 35-37 hCOs from 6 experiments (48 h experiments). Due to the nature of the original screen, control and nifedipine datapoints overlap with data published in Devilée et al. (Figure 1g and figure S1k) (39). *p < 0.05, ***p < 0.001, #p < 0.0001 using a Two-way ANOVA with Dunnett’s multiple comparisons test compared to DMSO at each time point (b-d) or a Kruskal-Wallis test with Dunn’s test for multiple comparisons (e-f) compared to DMSO. Data are presented as mean ± SEM with data points representing the average of all experiments (b-d) or individual hCOs (e-f). Experiments were performed using the HES3 and PB005.1 cell line with hCOs cultured using the DM protocol (67). *Con continuous, Rec recovery, Rep repeat, BL baseline, Nif nifedipine, Mav mavacamten*.

### Cardiomyocytes commit to the cell cycle within the first 4 h of treatment

We next determined the timeframe in which the cell cycle response was established. hCOs were treated for 4 h followed by 44 h of recovery, for 24 h followed by 24 h recovery or treated for 48 h (Figure 3a). When treatments were removed after 4 h, hCOs recovered full contractility after removal of mavacamten and to 39% of baseline force for nifedipine-treated hCOs within 20 h (Figure 3b). Contraction rate, 50% activation and relaxation time also recovered (Figure S5). This transient 4 h treatment increased the number of Ki-67 positive cardiomyocytes in both nifedipine and mavacamten treated hCOs at experimental endpoint (48 h), whereas at least 24 h of treatment with AZ3 was required to see an increasing trend (Figure 3c). The 4 h transient treatment with mavacamten led to commitment to the cell cycle and resulted in an increased cardiomyocyte nuclei number at 48 h, which did not occur in nifedipine treated hCOs (Figure 3d). This finding was confirmed using transient 4 h aficamten treatment (Figure S6). Together these results highlight that cell cycle blockade can be alleviated with an acute 4 h reduction in active force using mavacamten or aficamten.

**Figure 3.**
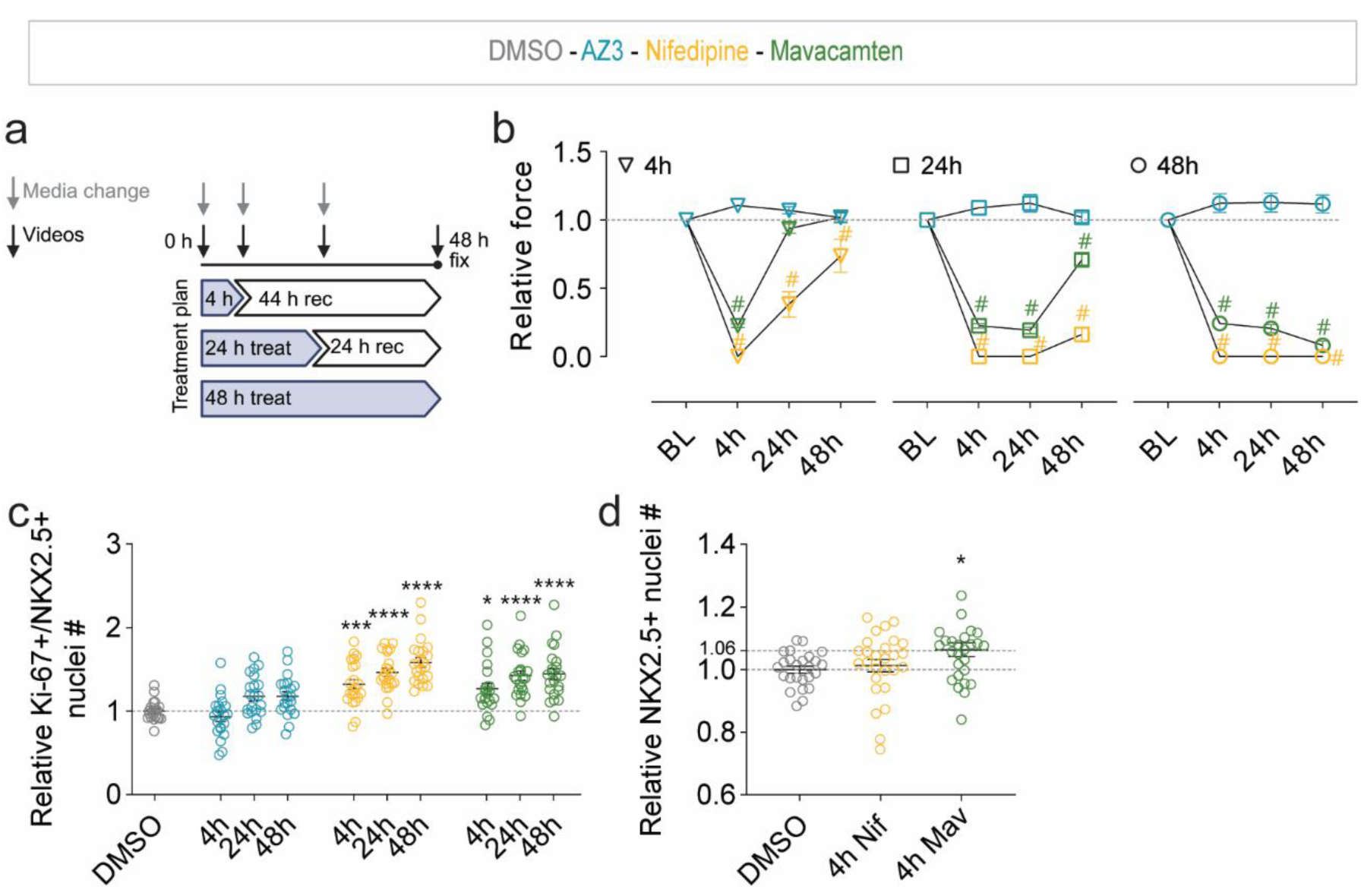
4 h functional inhibition was the critical time window during which cardiomyocytes can commit to cell cycle activity. (a) Schematic of the experiment. AZ3 was used at 3 µM, nifedipine at 3 µM and mavacamten at 0.3 µM. (b) hCO function over time displays functional recovery after removal of treatment compounds. N = 25-42 hCOs from 5 experiments. (c) Assessment of cell cycle activity at end point after short term treatments followed by recovery demonstrate 4 h of treatment is sufficient for cell cycle activation. N = 21-22 hCOs from 4 experiments. (d) Assessment of complete karyokinesis based on cardiomyocyte nuclei number after 4 h treatment. N = 25-29 hCOs from 5 experiments. *p < 0.05, ***p < 0.001, #/****p < 0.0001 using a Two-way ANOVA with Dunnett’s multiple comparisons test (B) or a Kruskal-Wallis test with Dunn’s test for multiple comparisons (c-e) compared to DMSO. Data are presented as mean ± SEM with data points representing the average of all experiments (b) or individual hCOs (c, d). Experiments were performed using the HES3 and PB005.1 cell line with hCOs cultured using the DM protocol (67). *BL baseline, Nif nifedipine, Mav mavacamten*.

### Phosphoproteomics reveals immediate changes in epigenetic and transcriptional control

Phosphoproteomics was performed to map the acute signaling changes initiated by active force reduction. This was performed 15 min after treatment where mavacamten and nifedipine reduced active force by 20% and ∼100%, respectively (Figure 4a-c). A total of 4046 unique phosphosites were identified on 1562 proteins. Nifedipine significantly changed phosphorylation on 112 sites while mavacamten affected 141 sites, with 68 phosphosites overlapping between the two groups (Figure 4d, Data S1). The direction of change for these overlapping phosphosites was consistent between nifedipine and mavacamten groups, indicating both treatments elicit similar acute signaling changes. The genes encoding the phosphoproteins were assessed on the presence of known single nucleotide polymorphisms (SNPs) in the genome that are causally associated with phenotypic cardiac traits. Heritability and posterior inclusion probability (PIP) analyses (49) were applied to cardiac MRI Genome-Wide Association Studies statistics (50). The gene list significantly enriched for left ventricular ejection fraction, wall thickness and volume traits when compared to all promotors (Figure S7a), indicating the proteins explain populational differences in cardiac function and structure. We further examined which genes are mostly likely causal of these traits (Figure S7b-d). ALPK3, LMNA, TTN, FOXK1, NACA and MYO18B were found to harbor PIP >0.6 SNPs for left ventricular wall thickness, which can be controlled by several factors including hypertrophy and proliferation.

**Figure 4.**
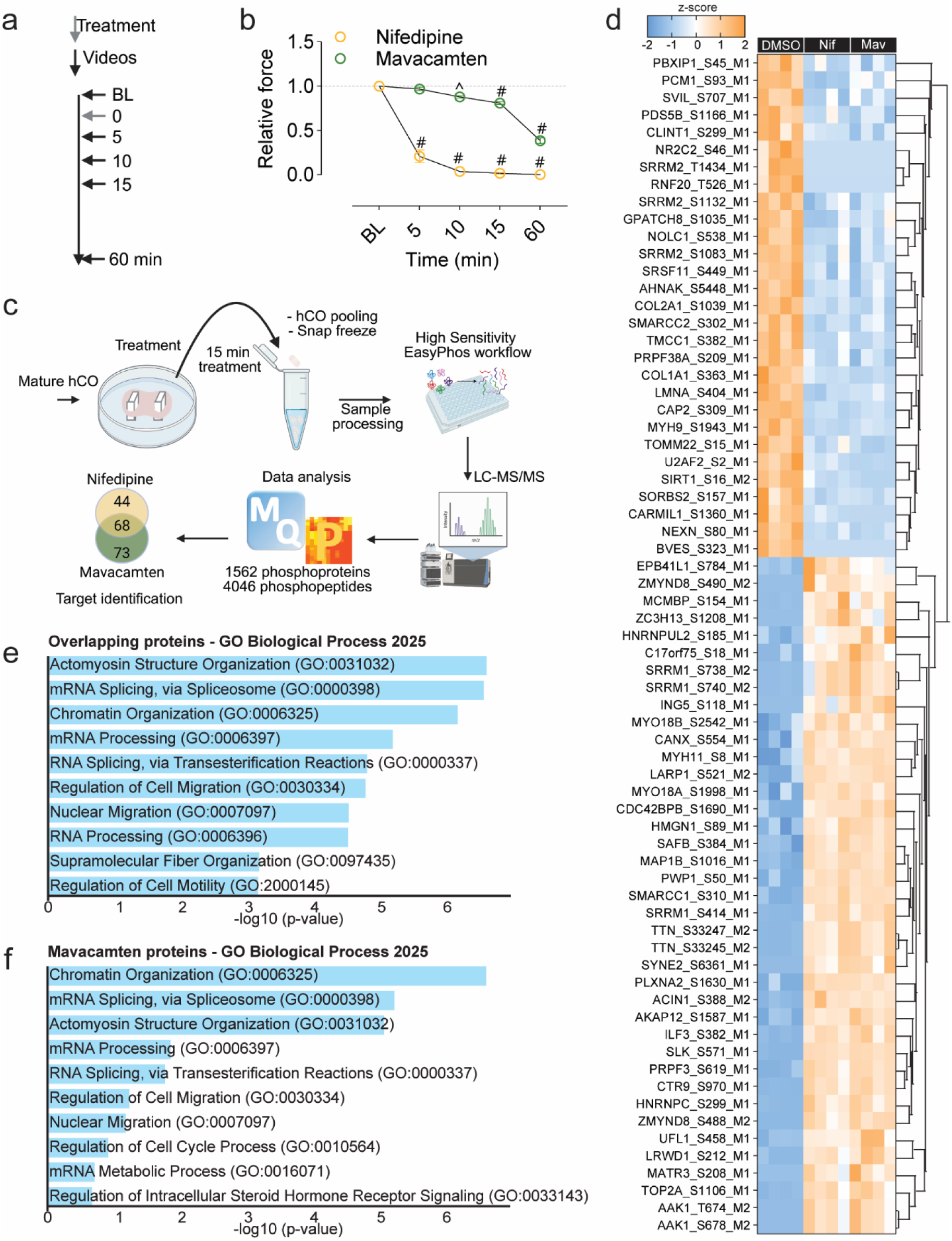
15 min functional inhibition impacts RNA processing, splicing and chromatin remodeling protein phosphorylation. (a) Schematic of the experiment for immediate functional assessment. (b) Functional response to nifedipine (3 µM) and mavacamten (0.3 µM) within the first 15 min of treatment up to 1 h of treatment demonstrating nifedipine and mavacamten induce a functional decline within minutes. N = 7-9 hCOs from 2 experiments. (c) Schematic of the phosphoproteomics workflow. hCOs were treated with nifedipine (3 µM), mavacamten (0.3 µM) or AZ3 (3 µM). A total of 4046 unique phosphopeptides were identified on 1562 unique phosphoproteins. (d) Heatmap displaying significantly affected overlapping phosphosites. N = 4 replicates per conditions, with each replicate consisting of 22-24 hCOs. (e) Gene ontology analysis for Biological Processes (2025) using Enrichr on the overlapping proteins associated with the phosphosites displayed in (d) highlights that these proteins enrich for RNA splicing and processing GO-terms. (f) Gene ontology analysis for Biological Processes (2025) using Enrichr on the mavacamten dataset demonstrates enrichment for proteins involved in cell cycle regulation. ^p <0.01, #p < 0.0001 using a two-way ANOVA with Dunnett’s multiple comparisons test compared to DMSO. Significantly changed phosphosites were identified based on a two-way ANOVA. Individual data points represent the average of the individual hCOs (b). Experiments were performed using the HES3 and PGP1 cell lines using the DM media protocol (67). *BL baseline, Nif Nifedipine, Mav mavacamten*.

Gene ontology analysis of the overlapping affected proteins particularly enriched for RNA processing and splicing terms (Figure 4e). Three of the affected RNA binding proteins in the overlap group, NOLC1, BCLAF2 and MATR3, and one specific to the mavacamten group (U2AF2) were reported to undergo strong changes in expression in postnatal day 1 (P1) and P7 versus P14 mouse hearts. These genes were highly expressed in the early developmental stages (P1 and P7) while expression was dramatically reduced by P14, suggesting their involvement in mediating cardiac maturation (51). Phosphorylation of S208 on MATR3 has moreover been reported to promote proliferation of neural stem cells while loss of phosphorylation induced differentiation (52). The proteins affected in mavacamten-treated hCOs enriched for ‘Regulation of Cell Cycle Process’ in addition to the enrichments found in the overlap group (Figure 4f). The proteins linked to this term include AHCTF1, BIN1, CETN2, SMARCC1, SMARCC2 and ARID1A. These last three are part of the BAF chromatin remodeling complex (53) which regulates cell fate decisions, including OSKM-mediated reprogramming and cardiogenesis (54–56). Furthermore, ARID1A promotes cardiomyocyte maturation and suppresses proliferation by mediating YAP activity (57). This indicates that epigenetic reprogramming may play a key role in activation of the cell cycle response.

Collectively, these phosphorylation changes suggest functional inhibition potentially initiates a dedifferentiation-like response through multiple mechanisms including chromatin landscape, transcriptional control and post-translational modifications on sarcomere proteins, leaving a lasting impact that persists after removal of mavacamten.

### Inhibition of contraction reversibly disrupts sarcomere gene expression and organization

As the phosphoproteomics indicated regulation of chromatin and transcriptional processes, RNA sequencing was used to determine transcriptional consequences 8 h after treatment compared to DMSO (Figure 5a). Mavacamten induced 107 differentially expressed genes (DEGs) (98 up, 9 down) which were fewer compared to nifedipine with 839 DEGs (298 up, 541 down) (39) (Figure 5b, c). There were only 9 genes upregulated in mavacamten-treated hCOs (*GPRIN3, ABAT, TKN2, GJA5, KY, PALMD, MAP2K6, KCNS2* and an uncharacterized gene) (Data S2). The downregulated DEGs were enriched for terms related to cardiac function, structure and development (Figure 5d, e). 103 out of 107 mavacamten DEGs overlapped with the nifedipine dataset with consistent direction of change, indicating that this subset is regulated by loss in contractile function in both treatments (Figure 5e, Data S2). Transcription of the overlapping DEGs was predicted to be controlled by SRF, NFE2L1 (NRF2) and PITX2 (Figure 5f), which are critical transcription factors for cardiomyocyte survival and regeneration *in vivo* (58, 59). As the transcriptional response was characterized by a reduction in the expression of sarcomere proteins (Figure 5g), immunostaining and analysis of organization (SOTA tool) (60) was used to profile the sarcomere structure (Figure 5h). We found that the structure was severely disrupted after both 4 and 48 h with nifedipine or mavacamten (Figures 5i-j), with further deterioration at 168 h continual treatment (Figure 5k). However, sarcomeres were able to reorganize and mature after treatment removal (Figure 5k, l). This indicates that active contraction directly controls sarcomere transcription and structure, and disruption may be alleviating a cell cycle blockade.

**Figure 5.**
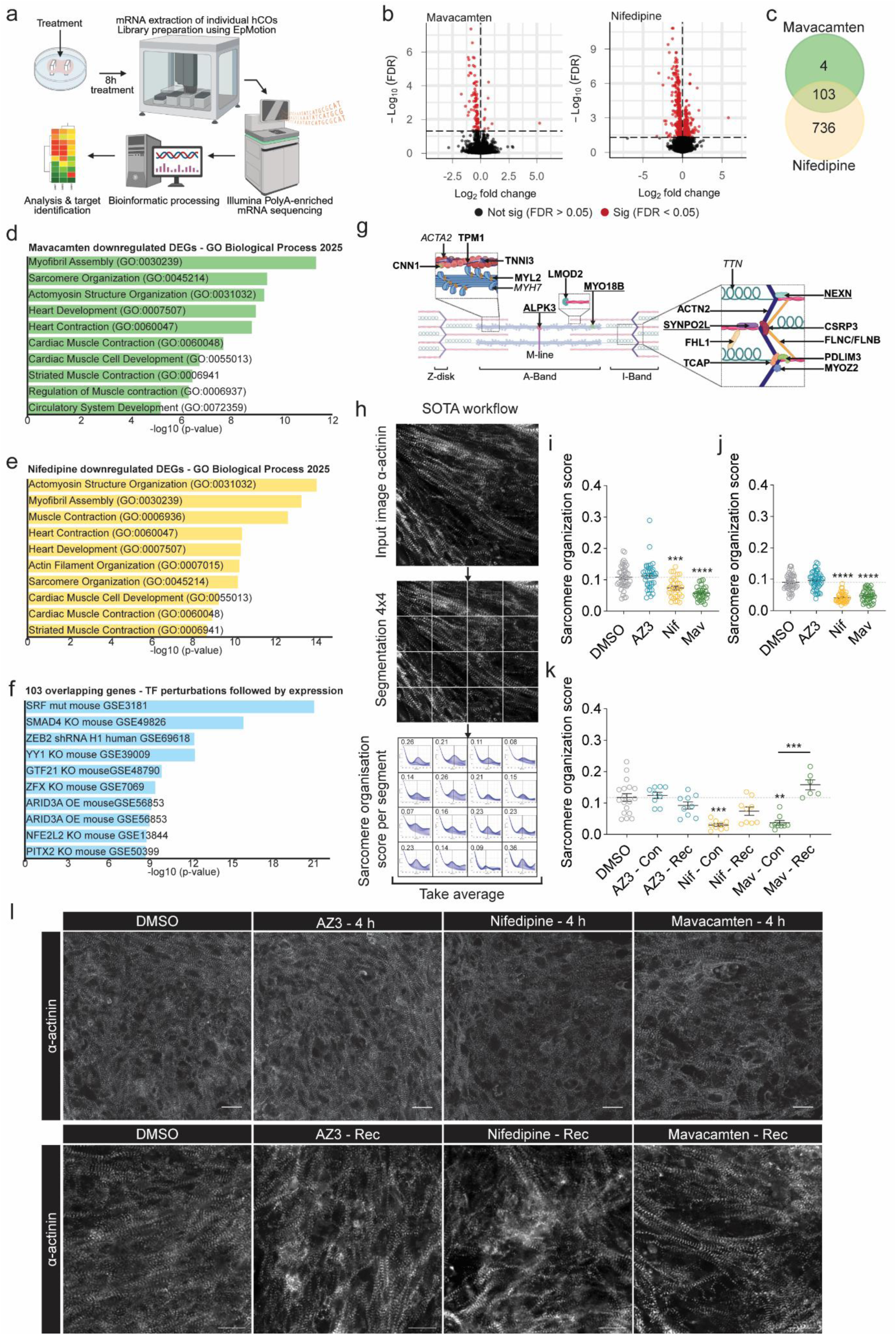
Sarcomeres are transcriptionally and morphologically disrupted following contraction inhibition. (a) Schematic displaying the RNA sequencing workflow. hCOs were treated for 8 h with nifedipine (3 µM) or mavacamten (0.3 µM) after which RNA was extracted from individual hCOs (N = 5 per conditions). DMSO treated hCOs were used as controls. (b) Volcano plot for differential expression analysis from comparisons mavacamten *vs* DMSO and nifedipine *vs* DMSO. Red dots indicate genes that are significantly regulated (FDR < 0.05). hCOs demonstrated a stronger transcriptional response to nifedipine with 839 differentially expressed genes (DEGs) while mavacamten differentially regulated 107 genes. (c) Venn diagram of statistically significant DEGs from comparisons mavacamten *vs* DMSO and nifedipine *vs* DMSO displaying a large overlap in transcriptional response. (d) Gene ontology terms (biological processes 2025) enriched for cardiac and muscle associated terms for genes downregulated by nifedipine. (e) Gene ontology analysis for biological processes (2025) for genes downregulated by mavacamten. (f) Gene ontology analysis for TF with perturbations for overlapping DEGs. (g) Schematic figure highlighting the downregulated sarcomere DEGs within the mavacamten RNA sequencing dataset. Genes in italic were not differentially expressed but are displayed to mark major sarcomere proteins. Genes encoding proteins that showed changes in phosphorylation are underlined. (h) Overview of the SarcOmere Texture Analysis (SOTA) workflow. An immunofluorescent image of α-actinin was used as input. Due to the 3D nature of the hCOs it was decided to perform image segmentation to improve analysis quality. For each image segment a sarcomere organization score was calculated. The average sarcomere organization score for each image was then used for analysis. (i) Nifedipine and mavacamten significantly reduced sarcomere organization score based on α-actinin remodeling at 4 h of treatment. N = 2 experiments, 30-45 images per condition. (j) Functional inhibition with either nifedipine or mavacamten for 48 h severely disrupts sarcomere organization based on a reduction in sarcomere organization score. N = 2 experiments, 39-48 images per condition. (k) Long-term treatment (168 h) further deteriorated sarcomere structure. Removal of treatment compounds (as per recovery treatment regimen (Figure 2a) allows for re-organization of the sarcomeres improving organization score. N = 1 experiment, 6-18 images per condition. (l) Representative images of α-actinin confirm visually disorganized sarcomeres in hCOs that were treated with nifedipine or mavacamten after 4 h of treatment. Sarcomere integrity improved after recovery from treatment. Scale bar = 20 µm. Differential gene expression analysis was conducted using gene-wise empirical Bayes quasi-likelihood F-tests for a given comparison. Benjamini-Hochberg method was applied on p-values for multiple testing correction. DEGs were identified based on an FDR <0.05. ***p< 0.001, ****p < 0.0001 using a Kruskal-Wallis test with Dunn’s test for multiple comparisons compared to DMSO (I,j) and between treatment pairs (k). Experiments were performed using the HES3 or PB005.1 cell line and cultured using the FBS (25, 68) (j) or DM (i, k) culture protocol (67). *Con continuous, Rec recovery, Rep repeat, Nif nifedipine, Mav mavacamten*.

### Contraction controls oxidative metabolism and aspartate/asparagine ratio

Metabolic dedifferentiation, characterized by the shift from oxidative metabolism towards the immature state characterized by more glycolysis is well-studied in proliferating cardiomyocytes (61–63). Oxidative metabolic activity was determined using a gas-trap design (Figure 6a) (64). The culture medium was supplemented with either ^14^C-palmitate or ^14^C-glucose and then ^14^C-CO_2_ was captured in the assay (Figure 6b). AZ3, which does not impact contraction, did not change oxidative metabolism, but both glucose and palmitate oxidation were substantially reduced (>50-70%) by nifedipine and mavacamten (Figure 6c, d). The glycolytic flux was determined by measuring ^13^C-lactate derived from exogenous ^13^C-glucose in the medium and was not impacted by any of the treatments (Figure 6e). This indicates that oxidative metabolism but not glycolysis is reduced by inhibition of contraction, and this may play a further role in the epigenetic regulation of cardiomyocyte proliferation (25, 65, 66).

**Figure 6.**
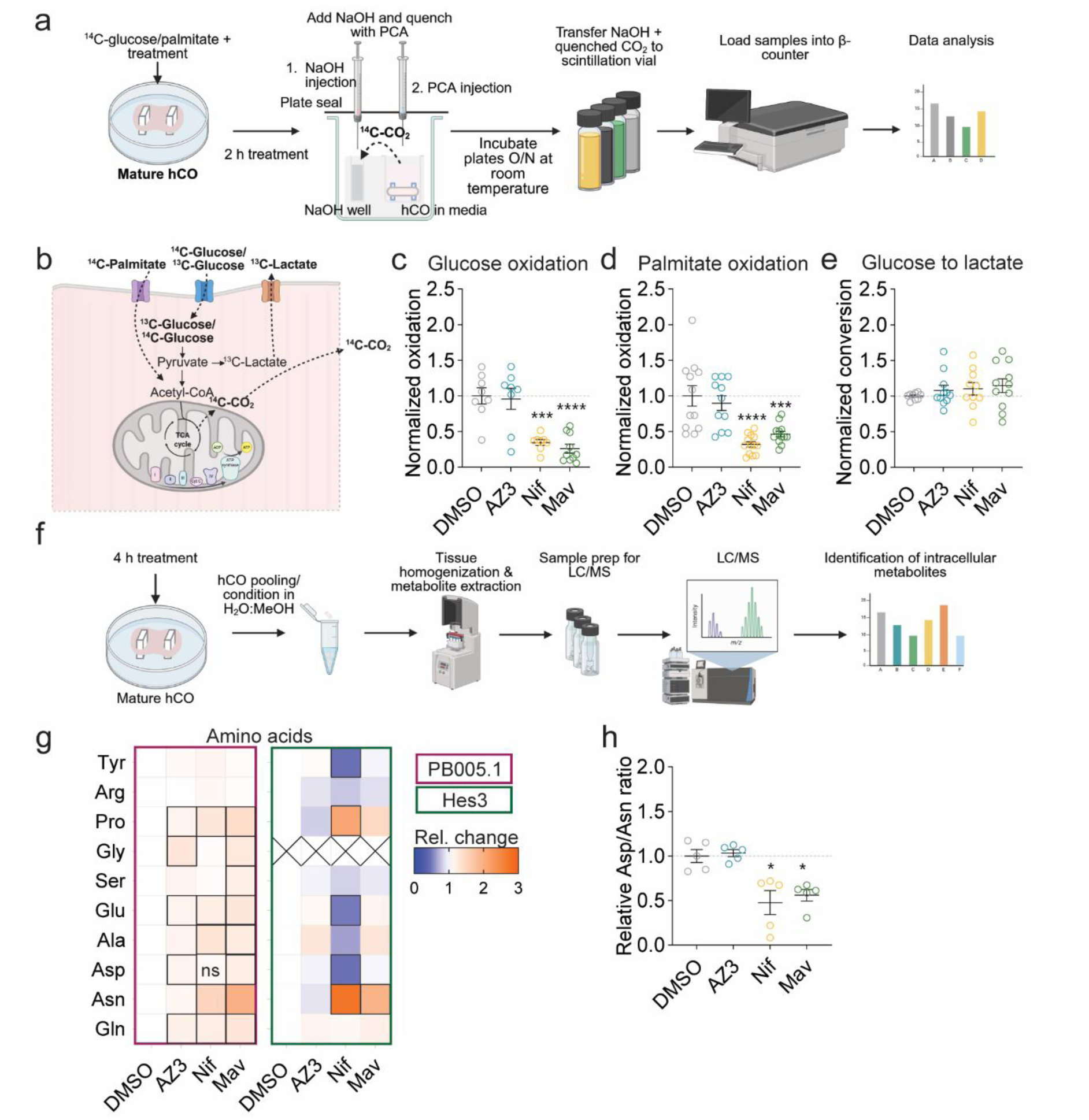
Metabolic rewiring following inhibition of cardiac function results shifts metabolism away from oxidative metabolism. (a) Schematic of the experiment with the workflow for the gas trap assay. hCOs were cultured in a custom-made PDMS well insert for a 12 well-plate. Each insert contained a well for the hCO and a well for NaOH. hCOs were treated with 3 µM AZ3, 3 µM nifedipine or 0.3 µM mavacamten. (b) Schematic of the metabolism of ^14^C-glucose/palmitate to CO_2_. (c) Nifedipine and mavacamten reduce glucose oxidation after 2 h of treatment. N = 8-10 hCOs from 2 experiments. (d) Inhibition of hCO function also reduced palmitate oxidation after 2 h of treatment. N = 11-13 hCOs from 3 experiments. (e) No changes were observed in the conversion of ^13C^-glucose to lactate. N = 9-11 hCOs from 3 experiments. (f) Schematic of intracellular amino acid extraction and identification using mass spectrometry. 6-8 hCOs were pooled per condition per replicate. (g) Heatmaps displaying identified intracellular metabolites after 4 h of treatment across 2 cell lines. Squares marked with a dark border depict statistically significantly changed metabolites. X marks missing values. N = 2 experiments with 2 cell lines, 2-3 replicates per condition per experiment. *p <0.05, ***p < 0.001, ****p < 0.0001 using a one-way ANOVA (c, d,e, h) with Dunnett’s multiple comparisons test compared to DMSO or a two-way ANOVA with Dunnett multiple comparison test (g). Data points represent individual hCOs (c, d, e, g). Experiments were performed using the HES3, PB005.1 and AA cell lines cultured using the FBS (25, 68) (c, d) or DM (e, g) culture protocol (67). *Nif nifedipine, Mav, mavacamten*.

A metabolic mediator, asparagine synthetase (Asns) was recently identified as a “universal” marker for cardiac dedifferentiation and regeneration (20). Asns converts aspartate (Asp) and glutamine (Gln) to glutamate (Glu) and asparagine (Asn). This is the only route through which Asn is synthesized *de novo*. To determine whether the amino acid profile of the hCOs was altered, intracellular metabolites were extracted and identified using LC/MS (Figure 6f). Asn levels were significantly increased across two cell lines and Asp to Asn ratio was decreased with 4 h of nifedipine or mavacamten treatment, indicative of rapid increased Asn synthesis and Asns activity (Figure 6g, h). These results indicate that active contraction also controls this metabolic marker of the dedifferentiation process.

## Discussion

While cardiomyocyte proliferation is associated with reduced contraction (19, 24–28), the cause and effect has not yet been explored. Here we used our hCO model (25, 26, 67, 68) to precisely study the impact of contraction on cardiac proliferation without the confounding hemodynamic effects *in vivo*. Targeting the myosin heavy chain using mavacamten enabled rapid control of active force production of hCOs without impacting other processes such as calcium cycling. Inhibition of contraction increased overall cell cycle activity, mitosis and cardiomyocyte nuclei number. Treatment with aficamten mirrored the mavacamten results, despite both drugs acting on different sites on the myosin heavy chain, providing further confidence for the specific involvement of contraction in blocking the cell cycle. Our findings fit with the observation that a degree of cell cycle reactivation occurs in mechanically unloaded hearts in patients where contractile demand is (partly) fulfilled by a mechanical pump (40, 41).

Signs of cell cycle activation were observed with 0.1 µM mavacamten, where contraction force was only reduced by 20% but the effects were stronger with 0.3 µM mavacamten. This highlights that active contraction does not require full inhibition, but at least a >20% reduction. Also, there was no benefit to long-term treatment with mavacamten or nifedipine. In contrast, a shorter treatment period was sufficient for cell cycle activation (nifedipine) and karyokinesis (mavacamten) despite recovery of function. This suggests responses are initiated within the first 4 h of treatment.

Our rapid small molecule approach enabled the study of mechanisms at very early timepoints through multi-omics techniques. The changes in phosphoproteome suggested an immediate impact on RNA processing and chromatin remodeling, which have been implicated in cardiac maturation and proliferation (69, 70). Moreover, there was a transcriptional loss of sarcomere and structural gene expression, which was confirmed by assessment of sarcomere organization depicting global sarcomere remodeling. These findings are consistent with reports of contraction controlling sarcomere maturation (71). Active remodeling of the sarcomeres is integrated in the sequence of events leading up to endogenous cardiomyocyte proliferation (8, 11, 72, 73). Co-immunoprecipitation of Tnnt2 one day after a myocardial infarction in P1 mice, a time at which sarcomere reorganization occurs, identified α/γ-adducin as potential mediators of this remodeling response (74). Follow up analysis demonstrated that adducin-induced sarcomere disassembly increased the number of proliferating cardiomyocytes, while mildly suppressing ejection fraction (74). Similarly, transiently forcing actin remodeling allowed for cardiomyocyte proliferation *in vitro* and *in vivo* in mice (75). The protein isoform composition of the sarcomeres is also important and changes dramatically during maturation. Troponin I undergoes isoform switching, from an immature skeletal muscle isoform (ssTnI, *TNNI1*) to a mature cardiac isoform (cTnI, *TNNI3*). This switch has been shown to control cardiomyocyte proliferation in mice (76). It is noteworthy that the regenerative zebrafish express a TnI isoform which lacks the mature cTnI N-terminus sequence thereby resembling the immature ssTnI (77). Consistent with this data, we observed a downregulation of the adult cardiac isoform (*TNNI3*) with both nifedipine and mavacamten treatment.

A shift to reduced oxidative metabolism and reliance on glycolytic activity has been shown to be critical to induce cardiomyocyte proliferation (16, 65, 78–80). Both nifedipine and mavacamten reduced oxidative metabolism to a similar level, highlighting that sarcomere contraction rather than calcium cycling uses the greatest amount of energy in the hCO contraction cycle. Our hCOs have a functional sarcoplasmic reticulum (67), so whether this is due to immaturity or the striking species differences in calcium handling between humans and other mammals and fish (81) should be explored by the field as this may have major implications for which processes to target to improve metabolism in heart failure (82). There was also a consistent shift in Asp/Asn ratio, indicative of increased Asns activity which has been identified as a “universal” marker of cardiomyocyte dedifferentiation and proliferation (20).

Collectively, we show that active contraction maintains the differentiated state of cardiomyocytes, including cell cycle block. That the cardiac maturation processes may be governed by active force rather than the other way around has key implications for the cardiac field.

## Methods

### hPS cell lines

Ethical approval for the generation and use of human heart tissue and hPS cells was obtained from QIMR Berghofer’s Ethics Committee and was carried out in accordance with the National Health and Medical Research Council of Australia regulations. Informed consent was obtained from all participants for the derivation and use of iPS cells including the PB005.1 (female) from MCRI. hPS cells were obtained from WiCell (Hes3, female), Coriell (PGP1) and the CIRM hPS cell repository (AA, male) funded by the California Institute of Regenerative Medicine. hPS cell lines are available upon request with appropriate agreements and ethical approvals.

hPS cell lines were maintained in mTeSR Plus (StemCell Technologies) in Matrigel-coated flasks (Millipore) and passaged using ReLeSR (StemCell Technologies). Karyotypes were routinely checked by G-banding (Sullivan Nicolaides) and molecular karyotyping analysis (Victorian Clinical Genetics Service and Ramaciotti Centre for Genomics). All cultures were routinely tested for mycoplasma. HES3 had a duplication at 20q20.11.

### Cardiac differentiation

A single-step differentiation protocol was used which generates ventricular cardiomyocytes and stromal cells simultaneously as previously described (25, 83). Four days prior (day −4) to the start of differentiation, hESCs/iPSCs were seeded onto Matrigel-coated flasks in mTesR Plus. Plating density was optimized for each cell line to obtain 50-60% confluence by day −1. Medium was changed to mTesR1 on day −1. Differentiation into mesoderm was initiated on day 0 by changing medium to RPMI 1640 GlutaMax (Thermo Fischer Scientific) (with 2% B27 minus insulin (Thermo Fisher Scientific), 200 µM L-ascorbic acid 2-phosphate sesquimagnesium salt hydrate (2AA, Sigma), 1% penicillin-streptomycin (P/S, Thermo Fischer Scientific)) supplemented with 1 µM CHIR99021 (Stem Cell Technologies), 5 ng/mL FGF2 (R&D Systems), 5 ng/mL BMP4 (Stem Cell Technologies), 9 ng/mL Activin A (R&D Systems). This medium was refreshed daily for 3 days. On day 3, medium was changed to RPMI 1640 GlutaMax (Thermo Fischer Scientific) (with 2% B27 minus insulin, 200 µM AA, 1% P/S) supplemented with 5 µM IWP4 (Stem Cell Technologies) to specify the culture towards cardiomyocytes and stromal cells. To prevent IWP4 crystal formation, it was added fresh to the medium and sterile filtered prior to adding it to the cells. Medium was changed on day 6 to this same medium containing 5 µM IWP4, except B27 minus insulin was replaced by B27 with insulin (Thermo Fischer Scientific). Medium was changed every 2-3 days until day 13 on which the same medium was added without IWP4. Cardiac cells were cultured in this medium until harvest on day 15.

Cardiac cells were harvested on day 15 using 0.2% collagenase type 1 (Sigma) in 20% fetal bovine serum (FBS, Thermo Fischer Scientific) and PBS (with Ca^2+^ and Mg^2+^) for 1 h at 37 °C with 5% CO_2_ in a humidified incubator. To improve detachment, flasks were tapped every 15-20 minutes. After 1 h, cells were washed off the flask and collected in PBS (minus Ca^2+^ and Mg^2+^) and centrifuged for 3 min at 300xg. Supernatant was removed and the cell pellet was resuspended in 0.25% Trypsin-EDTA (Thermo Fischer Scientific). The digest was inactivated using MEMα GlutaMAX (Thermo Fischer Scientific) supplemented with 10% FBS, 200 µM 2AA and 1% P/S. The cell suspension was filtered through a 100 µm cell strainer (BD Biosciences) to obtain a single cell suspension. Cell count was determined using a hemocytometer.

### Heart-Dyno fabrication

Cardiac organoids are generated and cultured in Heart-Dyno 96-well plate inserts. These are custom-made using polydimethylsiloxane (PDMS) and SYLGARD 184 Silicone Elastomer (CBC Australia) in a 6:1 ratio. This mixture was poured onto an aluminum mold (ANFF-SA), which was then degassed to remove air bubbles and baked 70°C to cure the PDMS. The PDMS sheet was removed from the mold after which plates were generated through either one of two methods. The earlier plates were made by hole-punching each individual insert from the PDMS sheet and gluing this into each well of a 96-well plate using uncured PDMS. The new method involved bonding of the complete PDMS sheet to the bottom of a bottomless 96-well plate (Greiner and Corning) using a chemical bonding protocol as previously described (84). Prior to use, plates were sterilized by immersion with 80% vol/vol ethanol for at least 2 hours. Ethanol was removed and plates were dried after which they were lidded and exposed to 2x 30 min ultraviolet cycles in the biosafety cabinet. Prior to plating, Heart-dyno inserts were coated with 1% bovine serum albumin (Sigma) in PBS for 1 h at room temperature.

### hCO production and maturation

hCOs were made using a previously described protocol (25, 67, 68). For a full 96-well plate, 6.6×10^^6^ cells in suspension were centrifuged at 300xg for 3 min to obtain a cell pellet. From here, two different media protocols were followed, called FBS for the protocol that incorporates FBS into the culture protocol (25) and a novel, more advanced ‘directed maturation’ (DM) protocol for a serum-free version of this protocol including additional maturation factors (67). The specific protocol used is described in each figure description. For the FBS protocol, the cell pellet was resuspended in MEMα GlutaMAX supplemented with 10% FBS, 200 µM 2AA and 1% P/S, 33 µg/mL aprotinin (Sigma or MedChemExpress) (seeding medium). The cells were placed on ice while the matrix mix was prepared. On ice, 3 mg/mL acid-solubilized collagen 1 (Devro) was salt balanced with 10x DMEM (Thermo Fischer Scientific) and pH neutralized with 0.1 M NaOH before adding 9% Matrigel (vol/vol). The cells were then incorporated into the mix and 3.5 µL was pipetted into each well of the earlier prepared 96-well plate (after BSA solution was removed). Throughout plating, the plate and cell-matrix mixture were kept on ice. The plating was done either manually one by one or in a semi-automated fashion using a VIAFLOW 96 (Integra) electronic pipetting system which plates all 96 hCOs simultaneously. Plate were centrifuged for 10-20 seconds at 100xg before incubating at 37 °C in a 5% CO_2_ humidified incubator for 30– 45 min to allow for gelation of the matrix. 150 µL seeding medium (37°C) was added to each well and plates were incubated for 2 days during which self-organization around the PDMS poles occurred. On day 17, medium was changed to maturation medium (MM) (25), consisting of DMEM, no glucose, no glutamine, no phenol red (Thermo Fisher Scientific) supplemented with 1× GlutaMAX (Thermo Fisher Scientific), 200 μM 2AA, 1% P/S, 4% vol/vol B27 without insulin, 33 μg/mL aprotinin, 100 μM palmitate (conjugated to bovine serum albumin in B27, Sigma) and 1 mM glucose (Sigma). MM was refreshed on day 20. On day 22, medium was changed to weaning medium (WM), which consisted of DMEM, no glucose, no glutamine, no phenol red supplemented with 1× GlutaMAX, 200 μM 2AA, 1% P/S, 4% vol/vol B27 without insulin, 33 μg/mL aprotinin, 10 μM palmitate (conjugated to bovine serum albumin in B27) and 5.5 mM glucose and 1 nM recombinant human insulin (Gibco). hCOs were cultured in WM until day 27 with media changes every 2-3 days. Experiments were started on day 27 using WM unless stated otherwise.

Medium changes were done manually or using the VIAFLO 96 (Integra) electronic pipetting systems. The DM protocol differed in that the feeding medium was composed of MEMα GlutaMAX supplemented with 200 µM 2AA and 1% P/S, 10 ng/mL bFGF (Sigma or MedChemExpress), 10 ng/mL PDGF-BB (Sigma or MedChemExpress), 4% B27 plus insulin (vol/vol) and 2 µM CHIR99021. Furthermore, DM-MM was additionally supplemented with 10 ng/mL bFGF and 10 ng/mL PDGF-BB. On day 24 and day 27, DM-WM was supplemented with 3 μM DY131 and 10 μM MK8722 (both MedChemExpress). Medium was changed on day 28 to normal WM and experiments were started on day 30 using normal WM unless stated otherwise.

Treatment compounds mavacamten (Selleckchem), nifedipine (Sigma), AZ3 (AstraZeneca), aficamten (Focus Bioscience) were reconstituted in DMSO, with DMSO used as a loading control across experiments. Treatment compounds were added to WM and a full media change (150 μL) was performed unless stated otherwise.

### Functional analysis

Videos were taken from hCOs under environmentally controlled conditions (37 °C with 5% CO_2_) using the Leica Thunder microscope (LASX software, version 5.3.0). 10 second videos of 500 frames were obtained at baseline and after different treatment durations as described in each figure description. Videos were analyzed using a custom-written MATLAB script (25) or using Tempo.ai analysis software (Dynomics) (85). Functional changes were determined based on the ratio of timepoint/baseline parameter and normalized to the average of control hCOs.

### Calcium amplitude assessment

hCOs were incubated with treatment medium including Fluo-4 AM (Thermo Fischer Scientific) for 60 min (37 °C with 5% CO_2_). The medium was then refreshed to treatment medium without Fluo-4 AM and hCOs were incubated for another 30 min prior to imaging. 10 second videos were taken using the Leica Thunder microscope (magnification 5x, exposure time 25 ms, LED 470 nM, filter 510). To analyze calcium amplitude, three regions of interest were marked on the tissue and F/F0 was determined using a custom-written MATLAB script.

### Immunohistochemistry and analysis

At experimental endpoint, hCOs were fixed in 1% paraformaldehyde for 1h at room temperature and washed with 1x PBS (3x). hCOs were incubated in blocking buffer (BB) consisting of 5% FBS, 0.2% Triton X-100 (Sigma) in 1x PBS for at least 4 h at 4°C on a rocker. hCOs were incubated with primary antibodies (Rabbit anti-Ki-67, Cell Signaling Technologies, D3B5, 1:400; Mouse anti-Ki-67, 8D5, Cell Signaling Technologies, 1:400; Mouse anti-phospho-histone H3 Ser10, Merck, 1:200, Mouse anti-α-actinin, Sigma, 1:1000; rabbit anti-NKX2-5, E1Y8H, Cell Signaling Technologies, 1:400) in BB overnight (45 μL/well) at 4°C on a rocker. The next day, hCOs were washed with BB 1x 5 min and 1x at least 2 h at 4°C on a rocker before incubating with secondary antibodies (Goat anti-mouse IgG1, goat-anti rabbit, Alexa Fluor 488, 555 or 633, 1:400) and Hoechst33342 (Sigma, 1:1000) in BB overnight 4°C on a rocker. The plate was protected from light. hCOs were then washed again 1x 5 min and 1x for at least 2 h in BB before imaging. hCOs were imaged while still in the Heart-Dyno plate in PBS or fructose-glycerol clearing solution (60% vol/vol glycerol (Sigma), 2.5 M fructose (Sigma)) (86).

hCOs were imaged using the Leica Thunder microscope at magnification 5x for whole hCO intensity analysis or 20x magnification for tile scan analysis. For initial screening purposes, whole hCO intensity was measured using a custom batch processing MATLAB script which enables background removal, image intensity measurement and export to Excel (Microsoft).

For tile scan analysis, 6 images per hCO were obtained at 20x magnification which were stitched with a 10% overlap. Images were loaded into the TissueGnostics StrataQuest (version 7.1.1.129) application which was designed for hCO image analysis. Images were cleaned to allow for individual nuclei detection based on Hoechst intensity. Cardiomyocytes were identified based on NKX2.5 nuclear expression. Cardiomyocytes in active cell cycle were identified based on double positive nuclei for NKX2-5 and Ki-67. Selection thresholds were set for all images per experiment, thereby eliminating observer bias.

To quantify the percentage of cardiomyocytes in mitosis or positive for Ki-67, hCOs were removed from the Heart-Dyno plate and mounted on slides in ProLong Glass mounting medium (Thermo Fischer Scientific). Confocal images (63x magnification) were obtained using either the Zeiss 780-NLO point scanning confocal microscope or the Leica Stellaris 5 confocal microscope. 3 random images per hCO were obtained. Cardiomyocytes were identified based on α-actinin expression. The number of cardiomyocyte nuclei was manually counted. Then, Ki-67 positive cardiomyocyte nuclei were counted to obtain a percentage for cardiomyocytes in active cell cycle. Cardiomyocytes in mitosis were identified based on the presence of a pHH3 positive nuclei overlapping with α-actinin remodeling. The numbers of three images per tissue were combined to calculate the percentages per hCO.

### Phosphoproteomics

Medium was changed using the VIAFLOW 96 (Integra) to treat all wells simultaneously and hCOs were incubated for 15 min at 37 °C, 5% CO_2._ hCOs were then washed with ice cold TBS (50 mM TRIS-HCl (pH 7.4), 150 mM NaCl) (100 μL per well, 3x). Plates were kept on ice throughout this procedure. While in ice cold TBS buffer, hCOs were removed from the Heart-Dyno plate and pooled per condition (22-24 hCOs per condition, 4 replicates). Samples were snap frozen and stored at −80 °C for LC-MS analysis. Samples were lysed in ice cold SDC buffer (4% (wt/vol) SDC and 100 mM Tris-HCl (pH 8.5)) and heat treated at 95 °C for 5 min. Sonication was performed and protein concentrations were determined by BCA assay. Equal protein concentrations were obtained through dilution in SDC lysis buffer. Samples were incubated with reduction-alkylation buffer (100 mM TCEP and 400 mM 2-chloroacetamide) for 5 min at 45 °C while shaking at 1500 rpm to reduce disulphide bonds and carbamidomethylate cysteine residues. Samples were digested overnight by Lys-C and Trypsin (1:100 ratio of enzyme-to-substrate) at 37 °C while shaking (1500 rpm). Enrichment of phosphopeptides and desalting was performed using the EasyPhos workflow as described (87). Phosphopeptides were vacuum dried at 45 °C, reconstituted in MS loading buffer (0.3% (vol/vol) TFA, 2% (vol/vol) acetonitrile), and separated by reversed-phase chromatography using a 40 cm column with an inner diameter of 75 µm packed in-house with C18 material (1.9 µm (Dr. Maisch)) and maintained at 60 °C using a column oven (Sonation, GmbH). Peptides were separated by linear gradients of buffer A (0.1% (vol/vol) formic acid) and buffer B (80% (vol/vol) ACN and 0.1% (vol/vol) formic acid) and analyzed by LC-MS/MS as described previously (88). Data analysis was performed as previously described (88), using Spectronaut (version 16.0.220606.53000) and searches performed against the Human UniProt Reference Proteome (March 2022 release) with default settings (precursor and protein Qvalue cutoffs 0.01, Qvalue filtering, MS2 quantification). ‘‘PTM localization’’ was turned on. The PTM ‘‘Probability cutoff’’ was set to 0 and localization filtering (min. 0.75 localization probability) was performed during downstream analysis. Spectronaut output tables were processed using the Peptide collapse (v1.4.2) plugin for Perseus (89). Phosphopeptides were included in statistical analyses if there were at least 3 valid values for at least one condition. Excel was used to perform T-tests after which multiple testing correction to control false discovery rate (FDR) was performed. https://maayanlab.cloud/Enrichr/ was used for GO-term analysis using the ‘GO Biological Process 2025’ database.

#### Posterior inclusion probability analysis

GWAS summary statistics were quality-controlled with HapMap3 LD reference for consistent SNP frequency, allele and sample sizes. Heritability enrichment and posterior inclusion probabilities (PIPs) were computed using SBayesRC with baseline v2.2 model (90) to estimate trait-causality for each SNP. Genes ±5kb windows in the mavacamten dataset, including FOXK1, were mapped to hg19. FOXK1, which mediates cardiac proliferation (91), was included in the analysis as it was phosphorylated at S299 in all 4 nifedipine and mavacamten samples and only in 1 control. They were then compared against SNPs in all promotors ±500bp provided in the baseline v2.2 model using one-tailed student’s t-test for statistical significance of heritability enrichment. Manhattan PIP plots were generated using Python seaborn and matplotlib packages.hg198.

### RNA sequencing

#### RNA extraction and library preparation

Following 8 h of treatment with treatment compounds or DMSO vehicle control (5 hCOs per condition), hCOs were washed with PBS, lysed and stored at −80 °C. NucleoMag96 RNA kit (Macherey-Nagel) was used for RNA extraction on the EpMotion 5075t. RNA quantification was performed using Qubit RNA High Sensitivity Assay (Invitrogen) and integrity was determined using the 4200 TapeStation with High Sensitivity RNA ScreenTape kit (Agilent). Samples were normalized to ≤100 ng RNA using nuclease free water in a total of 25 µL. The Illumina Stranded mRNA Preparation Ligation kit and the IDT for Illumina RNA UD Indexes Set A (Illumina) were used to prepare poly(a)-enriched RNA libraries. Libraries were quantified using the Quant-iT dsDNA broad range (BR) assay kit (Invitrogen). The quality of libraries was assessed using the 4200 TapeStation with the D1000 ScreenTape kit (Agilent). Single-end 100 bp sequencing was conducted on the NovaSeq 6000 (Illumina).

#### Read mapping and quantification

Basecalling and adapter trimming was completed on BaseSpace (Illumina). ‘FastQC’ (version 0.11.9) was run for quality control. Reads were aligned using ‘STAR’ (version 2.7.9a) (92) to the GRCh38 assembly with the gene, transcript, and exon features of the Ensembl (release 106) gene model. BAM files for each sample were merged using samtools (version 1.9) (93). RSEM (version 1.3.1) (94) was used to quantify expression. Duplicate reads were marked using Picard MarkDuplicates in the GATK package (version 4.2.4.1). ‘RNA-SeQC‘ (95) (version 2.4.2) was used to compute quality control metrics. Org.Hs.eg.db (version 3.14.0) was used to annotate gene biotypes, which were all kept for further analysis.

#### Differential expression analyses

Differential expression analysis was performed using edgeR’s quasi-likelihood pipeline (96–98) with filtered raw counts as input. The glmQLFit function from edgeR was used to fit a quasi-likelihood negative binomial generalized log-linear model to the read counts for each gene. The design matrix was defined by model.matrix(∼0 + Group), where Group represented DMSO, mavacamten and nifedipine. The glmQLFTest function from edgeR was used to conduct gene-wise empirical Bayes quasi-likelihood F-tests for a given contrast. Testing was performed for the following contrasts: mavacamten *vs* DMSO and nifedipine *vs* DMSO. Multiple testing correction was performed by applying the Benjamini-Hochberg (BH) method on the p-values, to control the FDR. DEGs were determined using the cut-off FDR < 0.05.

The original experiment included DMSO, nifedipine and mavacamten-treated hCOs. The nifedipine data compared to DMSO was analyzed separately and published in (39). Here, the complete dataset with nifedipine, mavacamten and DMSO-treated hCOs was re-analyzed.

### Metabolomics

#### Metabolite extraction from culture medium

hCOs were treated for 2-4 h with treatment compounds with or without [U-13C]-glucose. Medium of individual hCOs was collected, snap froze and stored at −80 °C until sample processing. Metabolite extraction was performed as previously described (99). Medium was mixed with 4 volumes of extraction buffer (1:1 vol/vol methanol and acetonitrile with 1 μM morpholineethanesulfonic acid (MES) (Sigma-Aldrich) as internal standard). Samples were vortexed and centrifuged for 20 min at 16.000xg at 4 °C. 80 μL of supernatant was transferred to HPLC vials and analyzed by LC-MS (100). Calibration standards for lactate were diluted into naïve medium and extracted as described.

#### Intracellular metabolite extraction

hCOs were treated for 4 h with compounds after which plates were placed on ice for 2 min before medium was removed and hCOs were washed with 5% (w/v) mannitol. Mannitol was removed and plates were placed on dry ice to snap freeze the hCOs. Plates were stored at −80 °C until sample processing. hCOs were lysed using ice cold lysis buffer (1:1 v/v H_2_O:MeOH with 1:10.000 internal standard (MES 0.2 mM)). hCOs were removed from the Heart-Dyno plate and pooled per condition (7-8 hCOs) into 500 μL lysis buffer. Sample was transferred to 0.5 mL PreCellys lysing vials (Lysko-A). hCOs and cell free control samples (lysis buffer only) were homogenized using the MiniLys system (Bertin Technologies) for 3×30 sec at max speed. 450 µL of sample was transferred to 1.5 mL Eppendorf tubes and 450 µL chloroform was added. After vortexing, samples were centrifuged for 20 min at 16.000xg at 4°C and 250 µL of aqueous phase was transferred to Eppendorf tubes. Samples were dried using GeneVac EZ-2 lyophilizer and pellet was resuspended in 50 µL ACN/H_2_O (1:1). To improve resuspension, samples were vortexed and incubated in a sonicating water bath. Once the pellet had been resuspended, samples were centrifuged for 15 min, 16.000xg at 16 °C. 40 µL of supernatant was transferred to LC vials and the samples were loaded onto the LC-MS (100).

#### LC-MS

Metabolites were identified by LC using a 1290 Infinity II pump (Agilent) with an InfinityLab Poroshell 120 HILIC-Z column (Agilent, 2.7 µm particle size, 2.1 mm internal diameter x 100 mm length. Before use, the column was deactivated with 0.5% (vol/vol) phosphoric acid according to manufacturer’s protocol. Metabolites were separated using buffer A, consisting of 80:20 (vol/vol) acetonitrile/water containing 10 mM ammonium acetate and 5 µM medronic acid (Sigma) to improve sensitivity and reduce peak-tailing (101) and buffer B, consisting of 10:90 (vol/vol) acetonitrile/water supplemented with 10 mM ammonium acetate and 5 µM medronic acid (100). These buffers were used as per the following gradients: start and 2 min, 100% buffer A (flow speed 250 µL/min); 12 and 14 min, 40% buffer B (flow speed 250 µL/min); 14.5 min and 18.4 min, 100% buffer A (flow speed 500 µL/min); 18.5 min, 100% buffer A (flow speed 250 µL/min). Autosampler temperature was 4 °C, column temperature was 30 °C, and injection volume was 3 µL. The Agilent 6470 QQQ was used for MS analysis set at gas temperature 200 °C, gas flow 11 L/min, nebulized pressure 40 psi, sheath gas temperature 400 °C, sheath gas flow 12 L/min, capillary voltage 3000V (positive and negative mode), nozzle voltage at 0 V for negative and 500 V for positive mode. Acquisition was performed with 10 ms dwell time. Data was analyzed using the Skyline Software (version 20.2.0.343-a7a9e8c4f) (102). Metabolite peaks were normalized to internal standards and data is presented relative to control.

### Tracing of radiolabeled metabolites

hCOs were cultured in a special 12-well plate setup with custom-made PDMS inserts to allow for gas-trapping (64). Prior to the start of the experiment with treatment compounds, hCOs were washed with WM. Medium was then changed to treatment medium supplemented with radiolabeled glucose ([U-14C]-glucose, final concentration 2 µCi/mL (PerkinElmer NEC042X001MC)) or palmitate ([1-14C]-palmitate, final concentration 0.1 µCi/mL (PerkinElmer NEC075H250UC)). Plates were sealed with a TopSeal-A Plus (PerkinElmer) and wells were equilibrated with 5% CO_2_ (64). A second seal was placed on top and plates were incubated for 2 h at 37 °C, 0% CO_2_. Subsequently, 100 µL 2 M NaOH was injected into its designated well through the plate seals and hCO metabolism was quenched using 50 µL 3 M perchloric acid which was injected into the medium. A third plate seal was used to close the injection site holes and plates were incubated overnight at room temperature to maximize CO_2_ capture. 100 µL NaOH was transferred to scintillation vials (6 mL, PerkinELmer) containing 3 mL Ultima Fold XR scintillant (PerkinElmer). Samples were gently mixed by inversion and radioactivity was measured by liquid scintillation counting using the TriCarb 4910TR (PerkinElmer). Signal was corrected for background signal as determined by cell free control samples consisting of naïve media with corresponding radiolabels.

### Statistical analysis

GraphPad Prism was used for statistical analyses unless stated otherwise. The Kolmogorov-Smirnov test for normality was conducted. Then, differences between the two groups were examined for statistical significance with unpaired Student t-tests or Mann-Whitney test. To compare data consisting of more than two groups, we performed one- or two-way ANOVA tests followed by Dunnett’s post-test multiple comparisons, a Kruskal-Wallis test with Dunn’s test for multiple comparisons or in case of uneven variances, a Brown-Forsythe and Welch ANOVA with Dunnett T3 test for multiple comparisons to get the corrected p-value. A value of p<0.05 was considered significant. Error bars represent standard error of the mean (SEM) unless specified otherwise. Datapoints represent individual hCOs or the average of all experiment as specified in the figure legends.

## Acknowledgements

The Heart-Dyno molds were fabricated using the NCRIS enabled Australian National Fabrication Facility – Queensland, New South Wales and South Australian Nodes. We especially thank S. Doe and M. Cherrill at ANFF-SA (Government of South Australia supported). We also thank T. Nguyen and N. Waterhouse for microscopy, B. Griffen for development of Tempo.ai, M. Hodson and M. Plan for processing metabolomics samples, and S. Rae for his insights. This research was supported by Snow Medical Research Foundation grant no. SMRF2019-060 to J.E.H., QIMR Berghofer to J.E.H., an investigator grant from the National Health and Medical Research Council of Australia (GNT2008376 to E.R.P.) and an QIMR Berghofer COVID grant to J.R.K, and a Diabetes Australia grant (Y23G-KRYJ) to J.R.K. The Novo Nordisk Foundation Center for Stem Cell Medicine is supported by Novo Nordisk Foundation grant NNF21CC0073729 to E.R.P. and R.J.M. The funders had no role in study design, data collection and analysis, decision to publish or preparation of the manuscript.

## Contributions

L.A.C.D. and J.E.H. conceived the project. L.A.C.D., J.D.R., J.R.K., H.R.R, S.J.H., M.L, R.L.F., and C.S.Y.C. contributed to the acquisition and/or analysis of data. L.A.C.D., J.R.K. C.S.Y.C., N.J.P., M.D., R.J.M. and J.E.H. contributed to the interpretation of data. Q-D.W., E.R.P. N.J.P., S.R.F., J.R.K. and J.E.H. contributed reagents and resources. L.A.C.D. and J.E.H. wrote the original manuscript. L.A.C.D., J.D.R., J.R.K., H.R.R., S.J.H., C.S.Y.C, M.L., R.L.F., Q-D.W., M.D., E.R.P., N.J.P., S.R.F., R.J.M. and J.E.H. revised the manuscript. Competing Interests E.R.P., R.J.M. and J.E.H. are co-inventors on a patent relating to the Heart-Dyno device and human cardiac organoid maturation used in this study (WO2018035574A1 filed by the University of Queensland) which is licensed to Dynomics. E.R.P., R.J.M. and J.E.H. are co-inventors on a patent for cardiac regeneration therapeutics (WO2020186283A1 filed by QIMR Berghofer). E.R.P. and J.E.H. are co-inventors on a patent for the serum-free conditions supporting the vascular population used in this study (WO2024016058 filed by MCRI and QIMR Berghofer). J.E.H. is co-inventor on licensed patents for cardiac differentiation and engineered heart muscle, some aspects of which are used in this study (WO2015040142A1 and WO2015025030A1), which are licensed to MyriaMed and Repairon. E.R.P., R.J.M. and J.E.H. are cofounders, scientific advisors, and stockholders in Dynomics. J.E.H. and R.J.M. are co-inventors on a provisional patent filed by QIMR Berghofer on the DM-hCO conditions (2024902826). Q-D.W. is an employee of AstraZeneca. The remaining authors declare no competing interests.

