## Supplementary Data for "Contractile function maintains cardiomyocyte differentiation and inhibits cell cycle activity"

### Supplementary figures

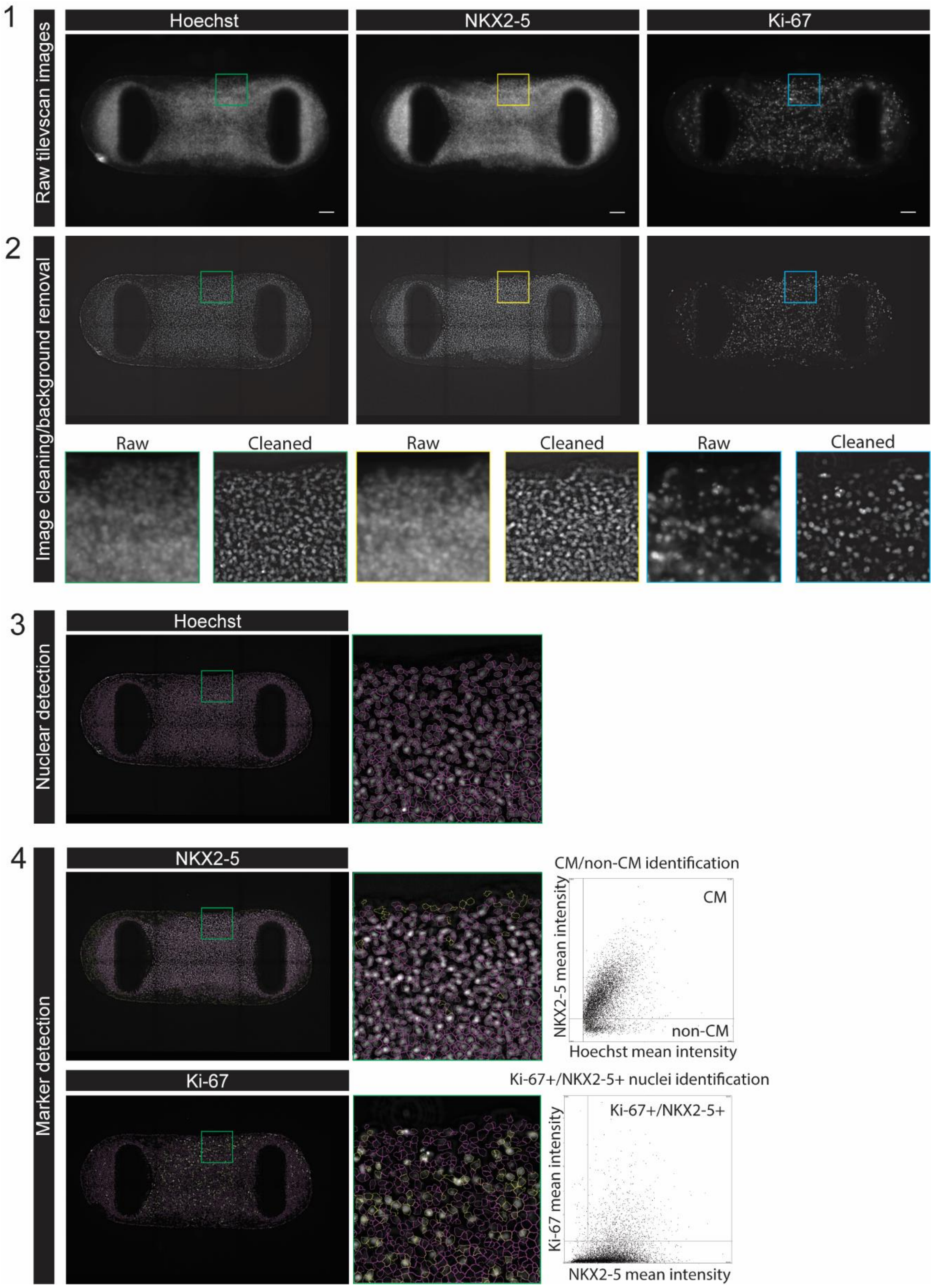

1
Raw tilevscan images
Hoechst
NKX2-5
Ki-67
2
Image cleaning/background removal
Raw
Cleaned
Raw
Cleaned
Raw
Cleaned
3
Nuclear detection
Hoechst
4
Marker detection
NKX2-5
CM/non-CM identification
CM
non-CM
NKX2-5 mean intensity
Hoechst mean intensity
Ki-67
Ki-67+/NKX2-5+ nuclei identification
Ki-67+/NKX2-5+
Ki-67 mean intensity
NKX2-5 mean intensity

**Figure S1. Overview of tile scan automated analysis for nuclear marker quantification.**
An overview of the different steps required to quantify the number of cardiomyocyte nuclei (NKX2-5 positive) positive for Ki-67. (1) 6 images (20x magnification) were obtained using the Leica Thunder microscope which were stitched to capture the complete hCO in one image. (2) Images were loaded into the TissueGnostics Strataquest application and image clearing was performed to remove background signal. A magnified view for each channel is provided before and after image clearing. (3) Nuclear segmentation was performed based on hoechst signal. Individual nuclei were segmented as shown in the magnified image. (4) The NKX2-5 nuclear intensity was used to set a threshold for NKX2-5 positive nuclei. The scatterplot demonstrates a clear separation of NKX2.5 positive and negative (non-cardiomyocyte) nuclei. Subsequently, a threshold was set to select for Ki-67 positive nuclei. The scatter plot demonstrates the Ki-67 positive cardiomyocyte nuclei in the upper right corner. The absolute numbers are exported into an excel sheet and used for further analysis.

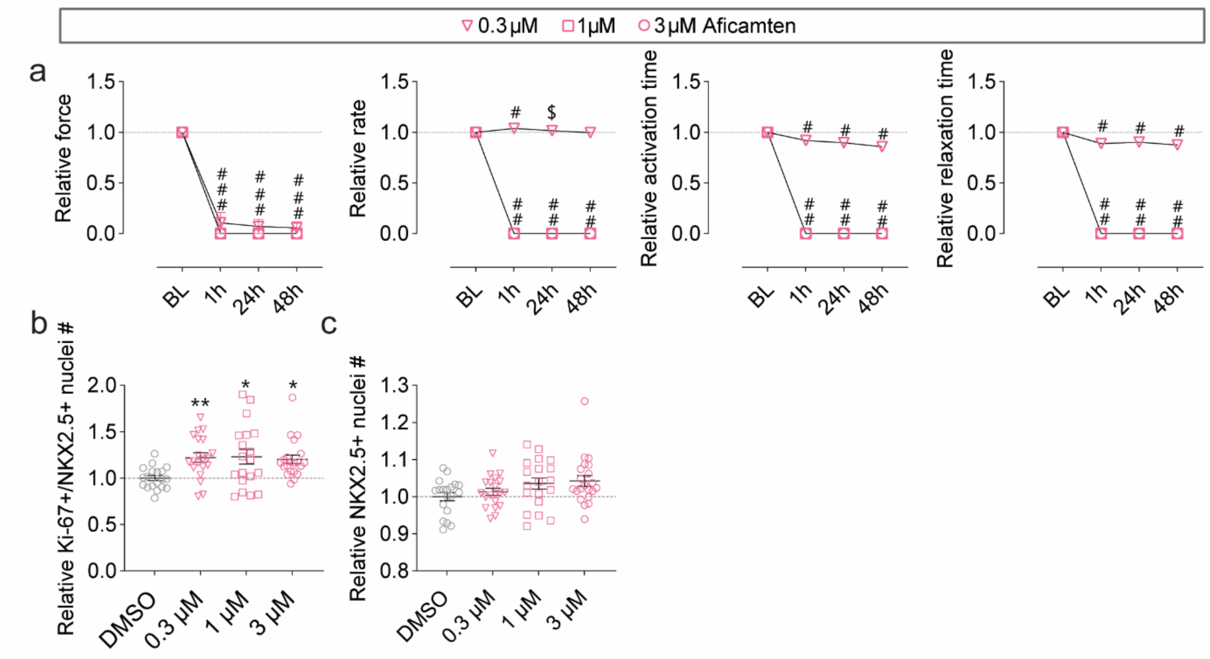


**Figure S2. Aficamten treatment leads to cardiomyocyte cell cycle activation.**

(a) Aficamten (0.3, 1 and 3 μM) induces an immediate and lasting functional response in hCOs. N = 21-22 hCOs per condition from 3 experiments.

(b) An increase in Ki-67 positive cardiomyocyte nuclei was observed across concentrations. N = 19-21 hCOs per condition from 3 experiments.

(c) An increasing trend in cardiomyocyte nuclei number was observed following 1 and 3 μM aficamten treatment for 48 h. N = 19-21 hCOs per condition from 3 experiments.

*p <0.05, ***/$p < 0.001, ****/#p < 0.0001 using a two-way ANOVA with Dunnett's multiple comparisons test compared to DMSO (a) or a Kruskal-Wallis test with Dunn's test for multiple comparisons (b, c). Significance is depicted for 0.3, 1 and 3 μM from top to bottom. Data is presented as mean ± SEM with data points presenting the average (a) or individual hCOs (b, c). Experiments were performed using the HES3 and PB005.1 cell lines cultured using the DM culture protocol (1). *BL baseline.*

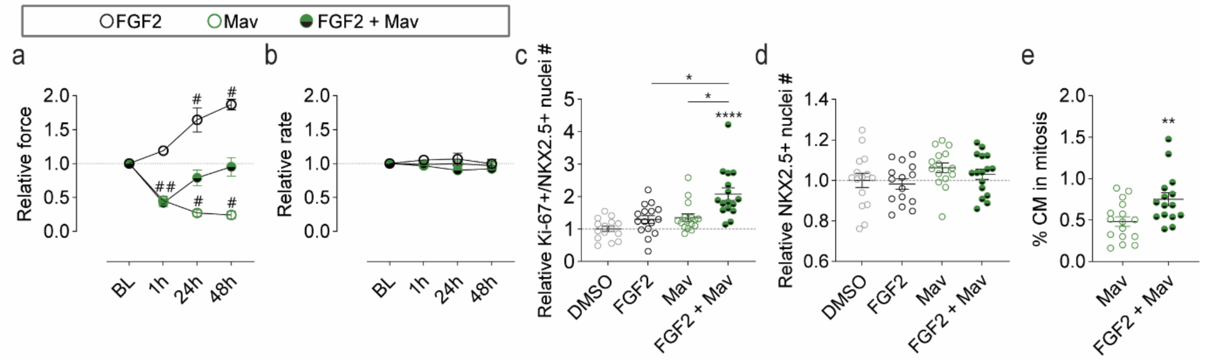


**Figure S3. Growth factor-boosted cell cycle activity may lead to mitotic catastrophe.**

(a) FGF2 treatment (10 ng/mL) increased force of contraction, even in combination with mavacamten (0.3 µM). N = 35-38 hCOs from 3 experiments.

(b) Rate was not impacted by FGF2. N = 35-38 hCOs from 3 experiments.

(c) FGF2 combined with mavacamten induced an additive effect on number of Ki-67 positive cardiomyocyte nuclei. N = 15-16 hCOs from 3 experiments.

(d) Combination treatment did not enhance cardiomyocyte nuclei number. N = 15-16 hCOs from 3 experiments.

(e) Manual quantification of the percentage of cardiomyocyte in mitosis demonstrated an additive effect between mavacamten and FGF2. N = 15-16 hCOs from 3 experiments.

*p < 0.05, **p < 0.01, #/****p < 0.0001 using a Two-way ANOVA with Dunnett's test for multiple comparisons (a, b), a Kruskal-Wallis test with Dunn's test for multiple comparisons (c), one-way ANOVA with Dunnett's test for multiple comparisons (d), or an unpaired t-test (e). Data are presented as mean ± SEM with individual data points representing average of all experiments (a, b) or individual hCOs (c-e). Experiments were performed using the HES3 cell line cultured using the FBS media protocol (2, 3). *BL baseline, Mav mavacamten.*

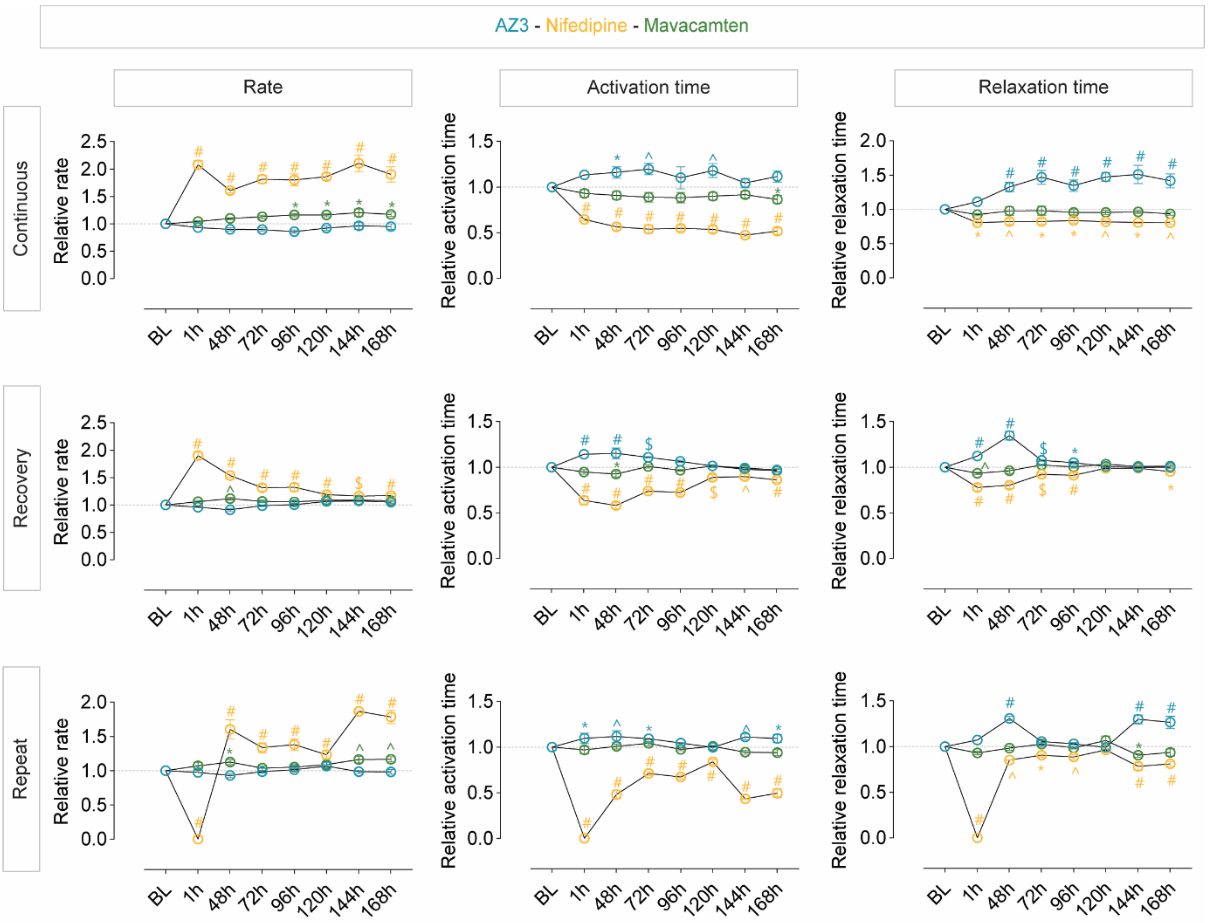


**Figure S4. Functional parameters following continuous treatment, recovery or repeat treatment.** Overview of the changes over time in rate, 50% activation and 50% relaxation times across treatment regimens as shown in Figure 2a. N = 4-6 experiments.

*p < 0.05, ^p<0.01, $p < 0.001, #p < 0.0001 using a Two-way ANOVA with Dunnett's multiple comparisons test compared to DMSO at each time point. Data are presented as mean ± SEM with data points representing the average of all experiments. Experiments were performed using the HES3 and PB005.1 cell line with hCOs cultured using the DM protocol (1). *BL baseline.*

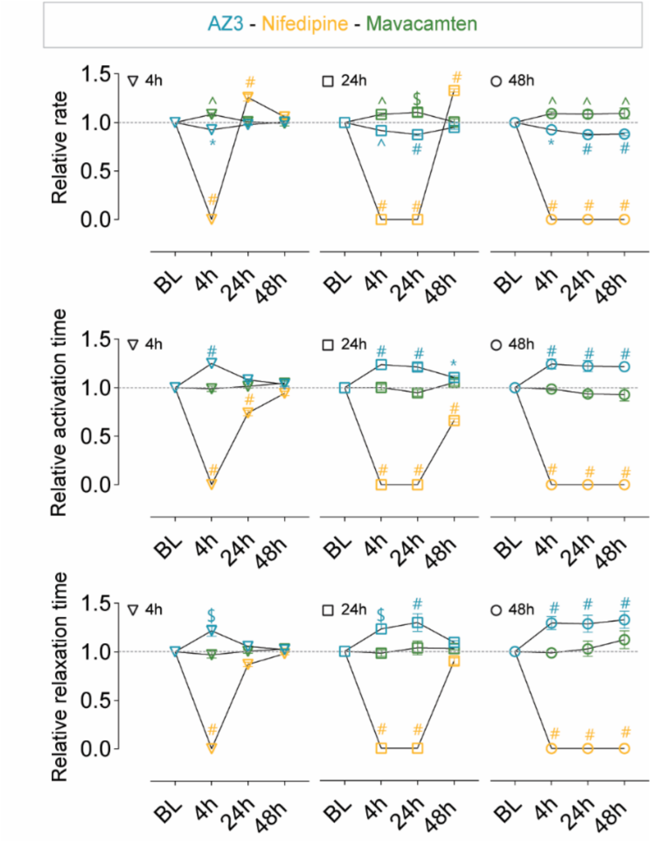


**Figure S5. Functional parameters recover following treatment removal.**

Overview of the changes over time in rate, 50% activation and 50% relaxation times upon removal of nifedipine (3 μM), mavacamten (0.3 μM) or AZ3 (3 μM) after 4 or 24 h of treatment as described in Figure 3a. N = 25-42 hCOs from 5 experiments.

* $p < 0.05$, ^$p<0.01$, \$$p < 0.001$, #$p < 0.0001$ using a Two-way ANOVA with Dunnett's multiple comparisons test compared to DMSO at each time point. Data are presented as mean ± SEM with data points representing the average of all experiments. Experiments were performed using the HES3 and PB005.1 cell line with hCOs cultured using the DM protocol (1). *BL baseline.*

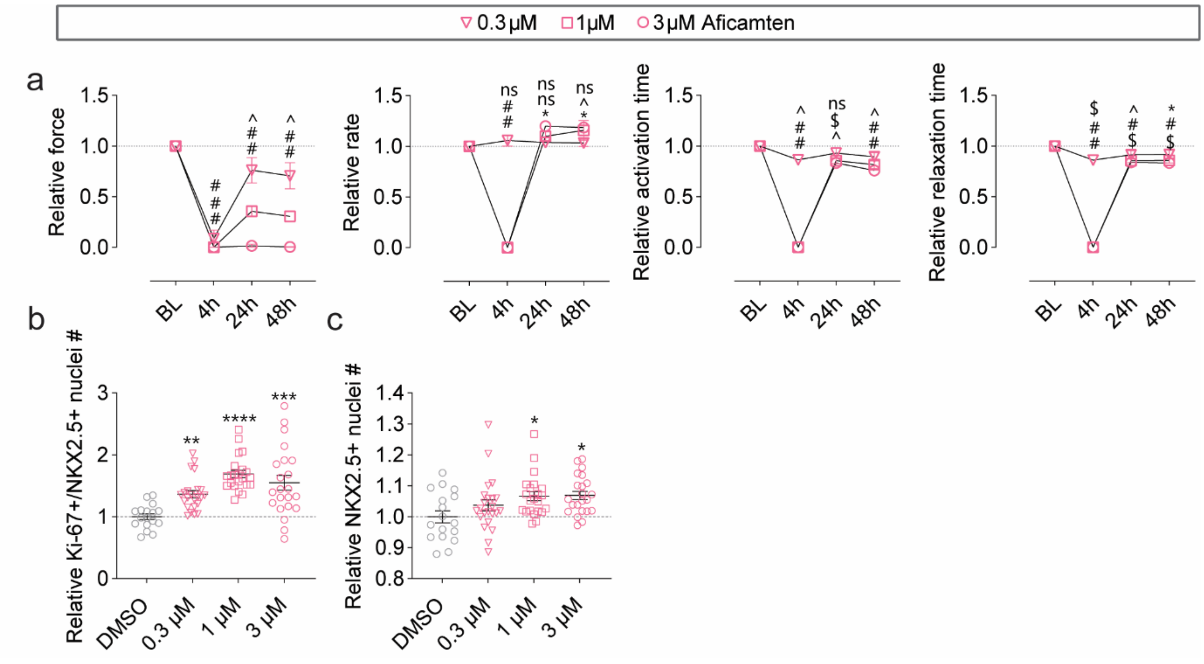


**Figure S6. Short-term reduction of active force production by aficamten initiates a cell cycle response.**

(a) Aficamten (0.3, 1 and 3 μM) reduces force of contraction with contraction partial recovery of force following treatment removal. N = 22-23 hCOs per condition from 3 experiments.

(b) An increase in Ki-67 positive cardiomyocyte nuclei was observed across concentrations. N = 17-23 hCOs per condition from 3 experiments.

(c) Short-term functional inhibition (4 h) is sufficient to increase cardiomyocyte karyokinesis based on an increase in cardiomyocyte nuclei number. N = 17-23 hCOs per condition from 3 experiments.

*p <0.05, ***/$p < 0.001, ****/#p < 0.0001 using a two-way ANOVA with Dunnett's multiple comparisons test compared to DMSO (a) or a Kruskal-Wallis test with Dunn's test for multiple comparisons (b, c). Significance is depicted for 0.3, 1 and 3 μM from top to bottom. Data points represent the average (a) or individual hCOs (b, c). Experiments were performed using the HES3 and PB005.1 cell lines cultured using the DM culture protocol (1). *BL baseline.*

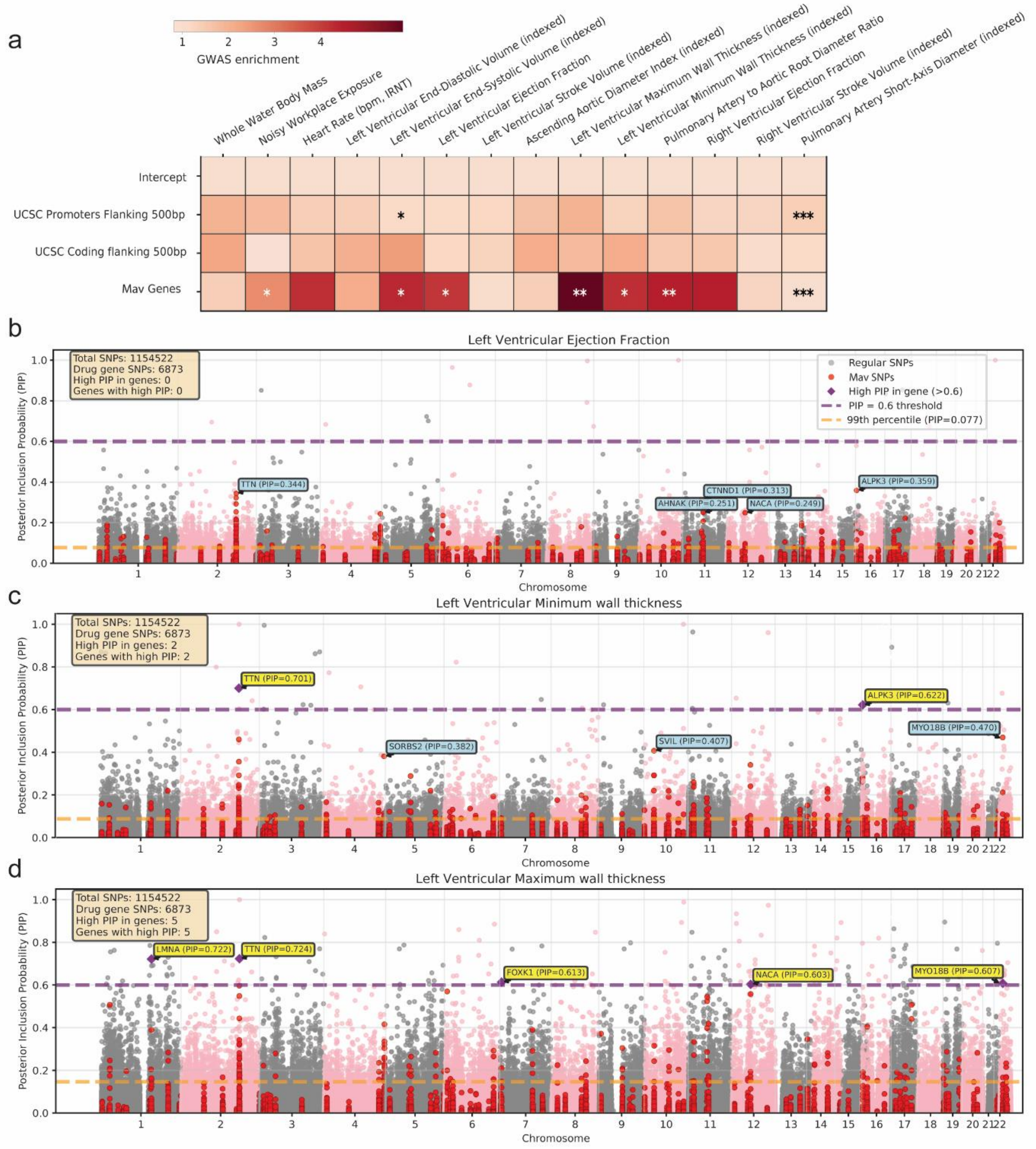


**Figure S7. Phosphorylated proteins contain common human genetic variation that explains population differences in cardiac function and structure.**

(a) Heatmap of drug-induced proteins show strong, significant enrichment for volume, ejection fraction, and wall thickness traits compared to all coding genes and promotors.

(b) Manhattan plot displays posterior inclusion probability (PIP) for 1 million SNPs for left ventricular ejection fraction. No genes meet the PIP threshold of 0.6 for causality. ALPK3, TTN, CTNND1, AHNAK, NACA are top 5 genes sorted by PIP.

(c) Manhattan plot for left ventricular minimum wall thickness. TTN, ALPK3 exceed promising likelihood of causality (PIP > 0.6), followed by MYO18B, SVIL, and SORBS2.

(d) Manhattan plot for left ventricular maximum wall thickness. TTN, LMNA exceed high likelihood for causality (PIP > 0.7), followed by FOXK1, MYO18B, NACA (PIP > 0.6).

Indexed means indexed to body surface area. *p <0.05, **p < 0.01, ***p < 0.001, using one-tailed z-test compared against enrichment of UCSC coding genes with flanking 500bp windows.
